# Reducing publication delay to improve the efficiency and impact of conservation science

**DOI:** 10.1101/2021.03.30.437223

**Authors:** Alec P. Christie, Thomas B. White, Philip Martin, Silviu O. Petrovan, Andrew J. Bladon, Andrew E. Bowkett, Nick A. Littlewood, Anne-Christine Mupepele, Ricardo Rocha, Katherine A. Sainsbury, Rebecca K. Smith, Nigel G. Taylor, William J. Sutherland

## Abstract

Evidence-based decision making is most effective with comprehensive access to scientific studies. If studies face delays or barriers to being published, the useful information they contain may not reach decision-makers in a timely manner. This represents a potential problem for mission-oriented disciplines where access to the latest data is paramount to ensure effective actions are deployed. We sought to analyse the severity of publication delay in conservation science — a field that requires urgent action to prevent the loss of biodiversity. We used the Conservation Evidence database to assess the length of publication delay (time from finishing data collection to publication) in the literature that tests the effectiveness of conservation interventions. From 7,415 peer-reviewed and non-peer-reviewed studies of conservation interventions published over eleven decades, we find that the mean publication delay (time from completing data collection to publication) was 3.6 years and varied by conservation subject — a smaller delay was observed for studies focussed on the management of captive animals. Publication delay was significantly smaller for studies in the non-journal literature (typically non-peer-reviewed) compared to studies published in scientific journals. Although we found publication delay has marginally increased over time (1912-2020), this change was weak post-1980s. Publication delay also varied inconsistently between studies on species with different IUCN Red List statuses and there was little evidence that studies on more threatened species were subject to a smaller delay. We discuss the possible drivers of publication delay and present suggestions for scientists, funders, publishers, and practitioners to reduce the time taken to publish studies. Although our recommendations are aimed at conservation science, they are highly relevant to other mission-driven disciplines where the rapid dissemination of scientific findings is important.

## Introduction

Across many mission-oriented disciplines, where there is an urgent need to tackle a societal issue, evidence-based decision making is critical to improving the effectiveness and efficiency of practice. This requires comprehensive access to scientific studies providing data useful for judging the likely effectiveness of actions. New scientific studies not only add to the relevant corpus of information that can guide decisions, but are likely to be particularly influential due to continually evolving technologies, methodologies, and skills, as well as the focus on issues of current concern. However, if new studies are not made available, or delayed in being so, relevant information useful for decision making (i.e., evidence; Salafsky et al. 2019) cannot easily be located by decision-makers in a timely manner. Not having rapid access to evidence to inform decision making risks suboptimal outcomes.

Biodiversity conservation is an example of a mission-oriented discipline and is motivated by a need for rapid, transformative change across the whole of society to tackle biodiversity loss (Mace et al. 2018; Leclère et al. 2020). Such an ambitious endeavour requires that we dramatically improve and upscale conservation efforts to reduce threats, and to protect and restore species and ecosystems. The urgency of the biodiversity crisis demands that conservation actions are as effective and efficient as possible, using the best evidence available to inform practice and policy (Sutherland et al. 2004; Pullin & Knight 2009).

Despite progress in the assessment of the effectiveness of conservation interventions (e.g., Sutherland et al. 2019), the evidence base is still patchy (Christie et al. 2020, 2021). Many commonly used interventions remain understudied and evidence for some threatened taxa or habitats remains non-existent or minimal for relevant conservation actions (e.g., Taylor et al. 2019; Junker et al. 2020). Without considering evidence on the effectiveness of actions, we risk implementing ineffective or potentially harmful actions and wasting limited conservation resources. Therefore, alongside the upscaling of conservation actions, we need more effective testing of interventions and streamlined channels to make evidence widely available.

When conservation interventions are tested, it is important to avoid unnecessary delays in publishing the results. Many issues in conservation are fast-moving and large delays could have detrimental impacts. For example, wind energy infrastructure has expanded massively around the globe from 489,000 MW in 2016 to 681,000 MW in 2019^1^, and delays to research papers testing cost-effective interventions to minimise bird and bat collisions could hamper key opportunities to mitigate the impacts of this expansion. In addition, one criterion for identifying Critically Endangered species is a ≥80% decline in population size over 10 years or three generations (IUCN 2019). In such cases, publication delay could occupy a substantial portion of the window for effective and efficient conservation action. Without well-targeted studies on species’ status, threats and responses, and timely publication of results, we risk mis-spending limited conservation funds on activities that are inefficient, ineffective, suboptimal or, at worst, harmful for biodiversity.

The problem of publication delay appears to be particularly acute in conservation. Kareiva et al. (2002) found that the mean time between the date of submission to the journal in which an article was eventually published (i.e., the destination journal) and publication was 572.2 days (1.6 years) in conservation science, far higher than for studies in genetics and evolutionary biology which had an average delay of 249.1 days (0.7 years). In 2009, a similar assessment looked at the same conservation journals, and found a destination journal delay of 402 days (1.1 years). This figure was still higher than for other biological fields (taxonomy = 334.5 days; behaviour = 379 days; evolution = 181 days), but had significantly declined over the previous seven years (O’Donnell et al. 2010). The same study investigated the delay between the completion of data collection and article submission, and found a median delay of 696 days (1.9 years), again higher than for the other fields studied (taxonomy = 605 days; behaviour = 507.5 days; evolution = 189 days; O’Donnell et al. 2010). If this same trend holds for papers that test the effectiveness of conservation interventions, a typical paper will take three years before it can help to inform the conservation community on the effectiveness of an intervention.

To examine the extent of this problem, specifically in the literature that tests conservation interventions, we investigate:

1. The length of publication delay in studies that test the effectiveness of conservation interventions, using the Conservation Evidence database (Sutherland et al. 2019).
2. How publication delay differs between different conservation subjects, publication sources (i.e., scientific journals or the non-journal literature), how this delay has changed over time, and how delay differs depending on the IUCN Red List (IUCN 2019) status of the species on which interventions are tested.

We define publication delay as the time taken from finishing the data collection for a study to when the study is published. We discuss the factors that could be driving publication delay and provide recommendations on how the scientific community can work together to minimise them.

## Methods

Using the Conservation Evidence database, we examined the difference between the year that data collection ended for a study and the year that the study was published. The Conservation Evidence database contains studies documenting 2,399 conservation interventions (as of December 2020; e.g., sowing strips of wildflower seeds on farmland to benefit birds) across multiple ‘synopses’. Synopses are used in the Conservation Evidence database to categorise studies into useful subject areas such as by species group, habitat, or related interventions (e.g., ‘Bird Conservation’ or ‘Management of Captive Animals’). To construct the database, publications were retrieved from the literature using a standardised protocol of manual searching through entire journals, and non-journal literature sources, for quantitative assessments of the effectiveness of a conservation intervention (‘subject-wide evidence synthesis’; see Sutherland et al. 2019 for details). For this analysis, we excluded reviews as we were interested in the publication delay of primary literature. We focused on the number of unique studies of an intervention within each Conservation Evidence synopsis. For example, if a publication reports studies of two different interventions (e.g., supplementary feeding and provision of artificial nests), then these studies are counted separately. Using this classification of conceptually distinct studies, we were able to extract information on when 7,415 studies were published and when their data collection ended. Approximately 3% of all studies did not report dates (280 out of 8,115 in the entire database as of December 2020) and so were excluded from the analyses.

Using the name of each study’s publication source and a dataset downloaded from SCImago (2020), we categorised the literature in which studies were published into three groups: i.) a ‘recognised journal’ (which had SCImago (2020) impact metrics — typically peer-reviewed journals); ii.) in ‘unrecognised journals’ (which did not have SCImago impact metrics — typically a mix of less conventional journals that may lack peer-review); and iii.) the ‘non-journal literature’ (often termed ‘grey literature’, which typically lacks peer review). This three-way separation of publication sources is a crude proxy for whether they are likely to be peer-reviewed (recognised journals = high; unrecognised journals = low-medium; non-journal literature = low) — thereby enabling some approximate estimation of the time taken for peer-review.

Where names of publication sources did not match the names given within the SCImago dataset, we manually searched to check whether names had changed over time, or had been incorrectly recorded in the Conservation Evidence database, and allocated publication sources to the ‘recognised journal’ category if a match was then found. Where names still did not match the SCImago dataset, we classified whether publication sources were either ‘unrecognised journals’ or from the ‘non-journal literature’ by manually reviewing the online information, such as the website, of the publication source. Any publication source – such as conference proceedings, books, theses, dissertations, reports, newsletters, and online articles – that did not make explicit mention of being a scientific journal was categorised in the ‘non-journal literature’ category. Any publication source that explicitly stated that it was a scientific journal was categorised as an ‘unrecognised journal’. For names of all publication sources in each of the three publication categories, see Tables S1–S3.

We extracted temporal information from the Conservation Evidence database (publication year) and a summary of each study that included information on when the study was conducted (the years the study began and ended). We defined the end year of a study as the year within which data collection ended (not when the intervention ended, for example). End years were extracted from Conservation Evidence summaries using regular expressions and text mining of the website (www.conservationevidence.com) with the XML package (Lang 2020a) and RCurl package (Lang 2020b) in R statistical software version 3.5.1 (R Core Team 2020). This extraction was necessary because this information is currently not in the database. We checked the accuracy of text mining by reviewing data for 79 studies (approximately 1% of the total number of studies analysed) and found that 94% had the correct study end year. Although there were a small number of errors, these were mostly caused by assigning the publication year as the study end year, and therefore would yield an underestimate of publication delay. In addition, automating the extraction of dates from study summaries offered the most feasible and reproducible way to analyse the entire evidence base, and avoided human error and unconscious bias that would affect manual extraction of dates (Christie et al. 2020, 2021).

To determine publication delay, we subtracted the end year of each study from its publication year. For studies conducted and published within the same year, their length of publication delay was therefore zero years. The coarse temporal resolution of years will have caused us to overestimate publication delay for studies with a delay of a few months which run between calendar years (e.g., December 2000 to March 2001), but underestimate the delay for studies published in months that do not span calendar years (e.g., January 2001 to December 2003). Across many studies these effects should generally cancel out – although rounding down of studies completed within one calendar year makes our overall estimations of publication delay conservative.

We used a Generalised Linear Model (GLM) to quantify and statistically test how publication delay varied: i.) between different synopses (Amphibian Conservation, Bat Conservation, Bee Conservation, Bird Conservation, Control of Freshwater Invasive Species, Farmland Conservation, Forest Conservation, Management of Captive Animals, Mediterranean Farmland, Natural Pest Control, Peatland Conservation, Primate Conservation, Shrubland and Heathland Conservation, Soil Fertility, Subtidal Benthic Invertebrate Conservation, Sustainable Aquaculture, Terrestrial Mammal Conservation); ii.) between different publication sources (recognised journals, unrecognised journals, and the non-journal literature; and iii.) over time (by publication year). For numbers of studies in each synopsis, please see Table S4. Therefore, we used three explanatory variables (synopsis, publication source, and publication year) to predict the response variable of publication delay. After an initial Poisson GLM revealed an overdispersion parameter value of 2.57, we used a quasi-Poisson GLM in which standard errors are corrected for overdispersion (using variance=theta*mu, where mu was the mean of the dependent variable distribution, and theta was the dispersion parameter of the quasi-Poisson model). The synopsis ‘Management of Captive Animals’ and publication source ‘non-journal literature’ were set as the intercept as these had the lowest mean publication delay values based on preliminary exploration of the data. We used Tukey’s all-pair comparisons (in the R package *multcomp;* Hothorn et al. 2008) to test for significant differences between synopses and between publication sources using our GLM. For the purposes of our visualisations, the plotted mean publication delay (and 95% Confidence Intervals) across all synopses was obtained using an intercept-only GLM. We used GLMs with a single fixed effect to plot summary statistics for visualisations of different synopses, different publication sources, and changes over time (i.e., using a fixed effect of either synopsis, publication source, or publication year, respectively).

As more recently published studies may be more likely to suffer from a longer delay, we conducted a sensitivity analysis by restricting the data we analysed in our original GLM to 1980-2020, 1990-2020, and 2000-2020 (Table S8) — we discuss these results later in the Results and Discussion.

In a separate analysis, we tested for significant differences in publication delay between studies testing interventions on species with different IUCN Red List statuses. To do this we extracted data from the Conservation Evidence database on the species studied within taxa-specific synopses (Amphibian Conservation, Bird Conservation, Terrestrial Mammal Conservation, Primate Conservation, and Bat Conservation), and the threat status of each species from the IUCN Red List (IUCN 2019). We limited the analysis to these synopses as these taxa had been comprehensively assessed in the IUCN Red List. We ran separate quasi-Poisson GLMs (using the same fixed effects of publication source, synopsis, and publication year, plus IUCN Red List Status) on a reduced dataset including all taxa-specific synopses, and on separate datasets for each of the taxa-specific synopses (Amphibians = Amphibian Conservation; Birds = Bird Conservation; Mammals = Terrestrial Mammal Conservation, Primate Conservation, and Bat Conservation).

For these taxonomic GLMs, we only considered the most threatened IUCN Red List category (out of Least Concern, Near Threatened, Vulnerable, Endangered, Critically Endangered) of all species for each published study. For example, if a study targeted multiple species, such as two that were listed as Least Concern and one listed as Endangered, we considered that as a study on an Endangered species. There were insufficient studies on species with IUCN Red List statuses of Data Deficient (zero studies) or Extinct in the Wild (less than eight studies) and so we did not include these categories in our taxonomic analyses. We were unable to obtain the IUCN Red List status of species at the time when the study was conducted and therefore had to use the current status of species in the latest IUCN (2020) Red List update. Whilst this may mean that, for some studies, certain species may have changed in their Red List status in the intervening years, the current threat category is an indication of the need for previous studies on responses that could have helped prevent this decline assuming that, for many species, threatening processes have been present over long time-periods. R code to perform all analyses is available here: https://doi.org/10.5281/zenodo.4621310.

## Results

The mean publication delay of studies of conservation interventions across all Conservation Evidence synopses was 3.57 years (95% Confidence Intervals = [3.50,3.64]; Fig. 1). Publication delay varied significantly between several synopses (p<0.05; Table S5-S6). Most notably, management of Captive Animals had many studies published in the same year as the end of the study, and a significantly smaller mean delay (2.0 years; Table S4; Fig.1) than most other synopses (p<0.01; Table S6) — except for Amphibian Conservation, Bat Conservation, Bee Conservation, Control of Freshwater Invasive Species, Primate Conservation, and Sustainable Aquaculture synopses (p>0.05; Table S6), each of which had a significantly smaller (p<0.05; Table S6) mean delay (<=3 years; Table S4) than most of the remaining synopses (each with a mean delay >3.5 years; Table S4; Fig.1).

**Figure 1.**
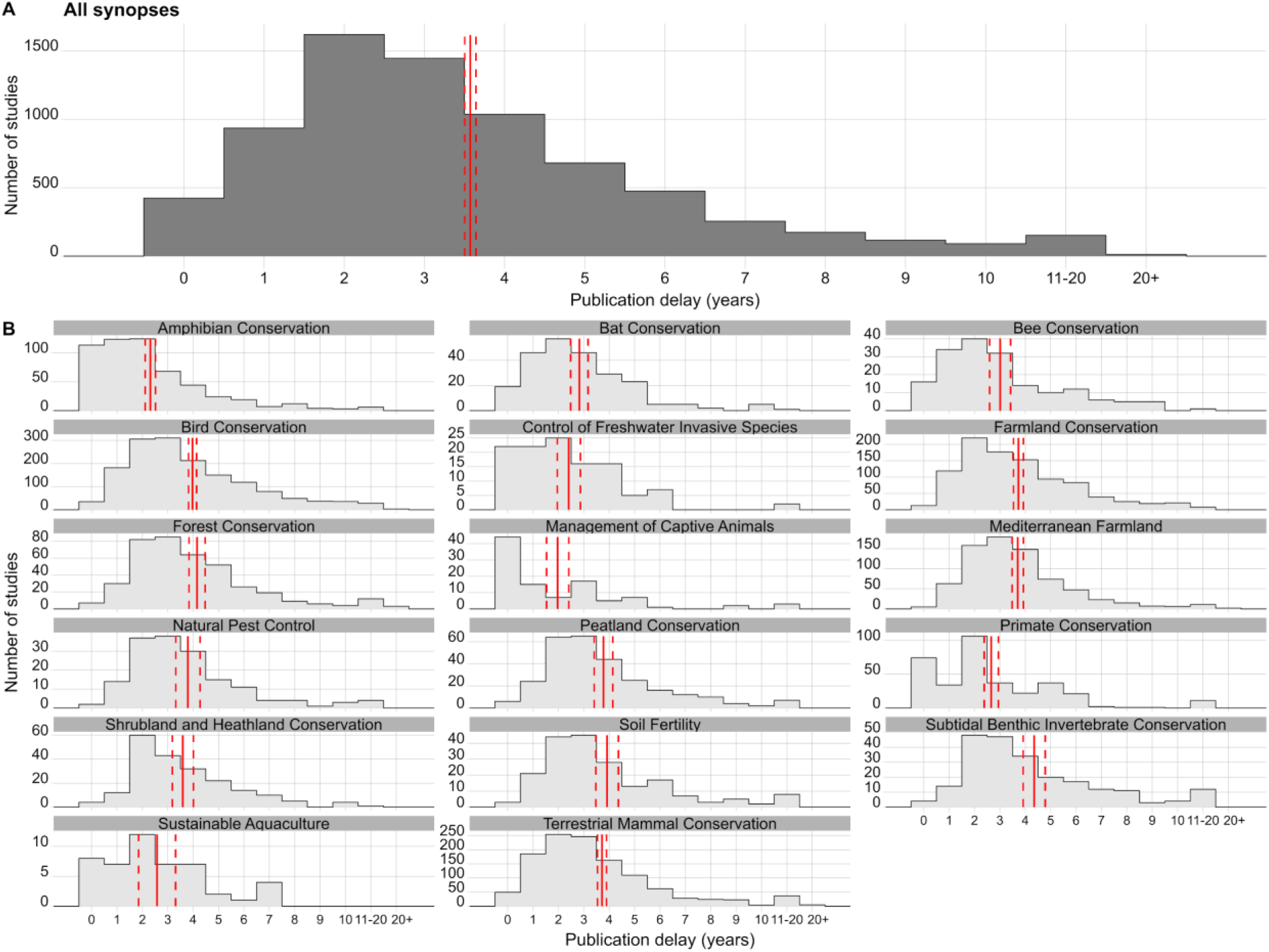
Distribution of studies of conservation interventions according to the length of publication delay (in years) for different Conservation Evidence synopses (each covering a distinct conservation subject — e.g., ‘Bird Conservation’). Solid red vertical lines indicate the mean length of publication delay for each plot and dashed red lines represent 95% Confidence Intervals. For each synopsis, summary estimates were obtained from a Generalised Linear Model (GLM) with only synopsis as a fixed effect, while estimates for all synopses were obtained from an intercept-only GLM).

Publication delay differed significantly by publication source (p<0.01; Table S5; Table S7; Fig.2); studies from non-journal literature (mean delay of 2.24; 95% Confidence Intervals = [2.06, 2.42]) had a significantly smaller delay compared to studies published in recognised journals (mean delay of 3.74; 95% Confidence Intervals = [3.66, 3.82]; t=9.9; p<0.001; Table S7). Studies from non-journal literature also had a significantly smaller delay than studies from unrecognised journals (mean delay of 2.73; 95% Confidence Intervals = [2.39, 3.08]; t=3.212; p=0.003; Table S7). Studies from recognised journals had a significantly greater delay than studies from unrecognised journals (t=-2.940; p=0.012; Table S7).

**Figure 2.**
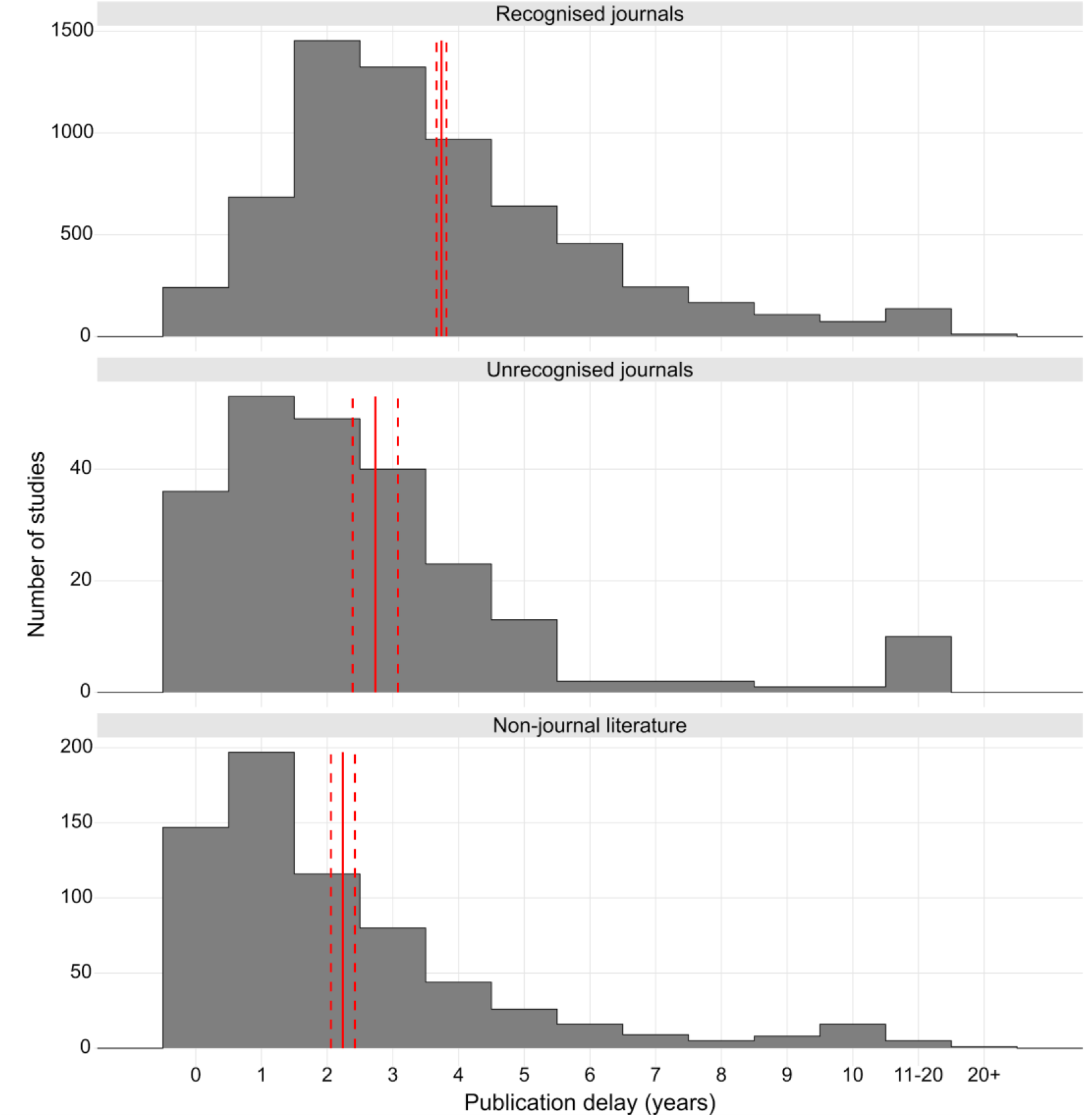
Publication delay in years for all studies of interventions published in recognised journals, unrecognised journals (according to SCImago (2020)), and the non-journal literature. Solid red vertical lines indicate the average length of publication delay for publication source and dashed red lines represent 95% Confidence Intervals. For each synopsis, summary estimates were obtained from a Generalised Linear Model (GLM) with only publication source as a fixed effect. For journals classified under each of the three categories see Tables S1-S3.

There was a small, but statistically significant, increase in publication delay from 1912 to 2020 (Fig. 3; t=3.598; p<0.001; Table S5); based on sensitivity analyses, a statistically significant increase in publication delay was also observed since 1980 (t=2.2; p=0.026; Table S8), but not since 1990 (t=0.3; p=0.787; Table S8) or 2000 (t=−0.13; p=0.192; Table S8).

**Figure 3.**
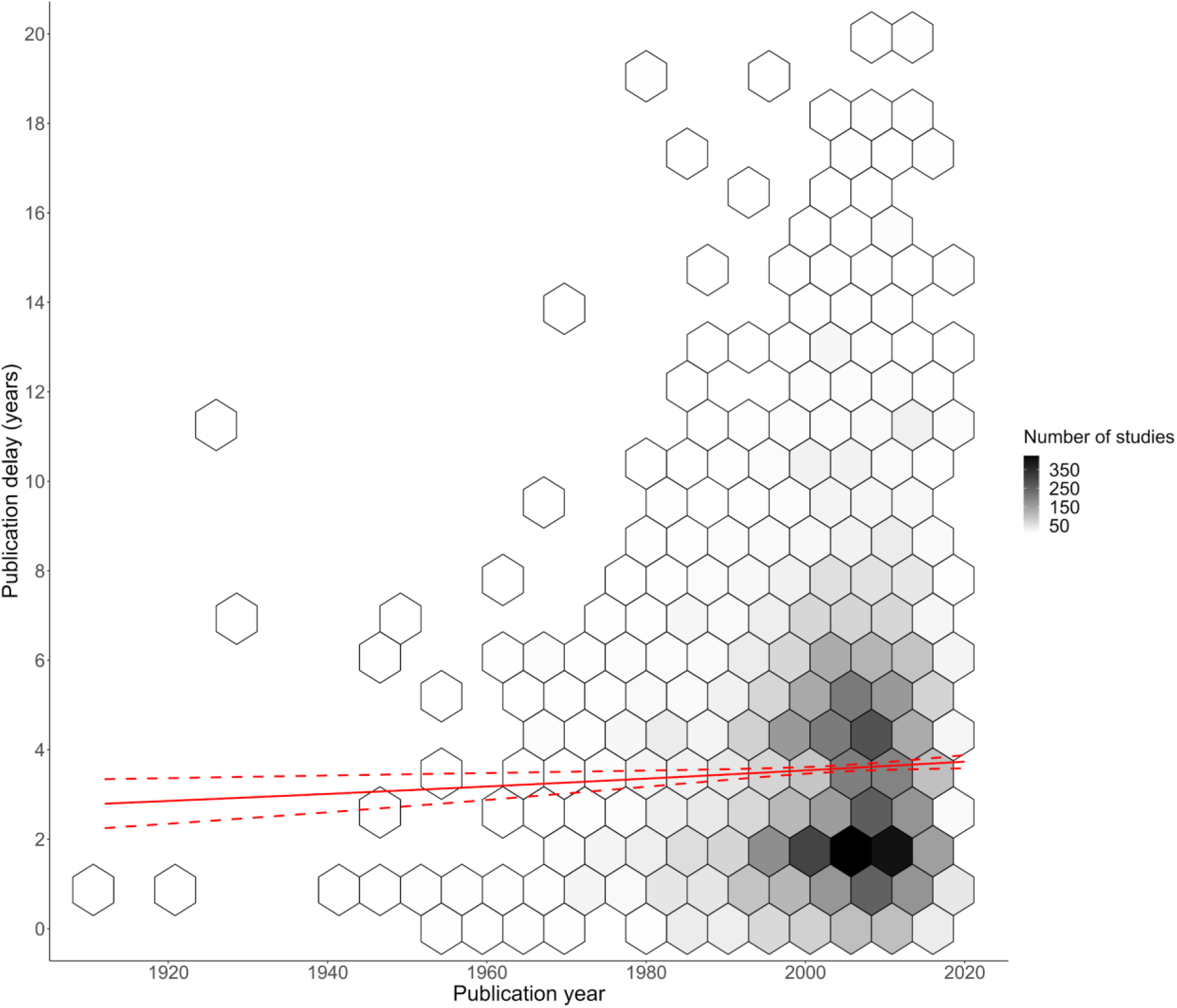
Changes in publication delay relative to the year in which studies of interventions were published. The shade of hexagons is relative to the number of data points (studies) at that position on the graph. The red solid and dotted lines represent modelled mean and 95% confidence intervals for publication delay using a quasi-Poisson Generalised Linear Mixed Model (GLM) (with only publication year as a fixed effect; see Table S5 for full model result). Only data for a publication delay of 20 years or less is presented to improve visualisation, but all data were used in the GLM (see full data figure Fig.S1). We conducted sensitivity analyses to check whether the trend changed in more recent decades (see Table S8).

**Figure 4.**
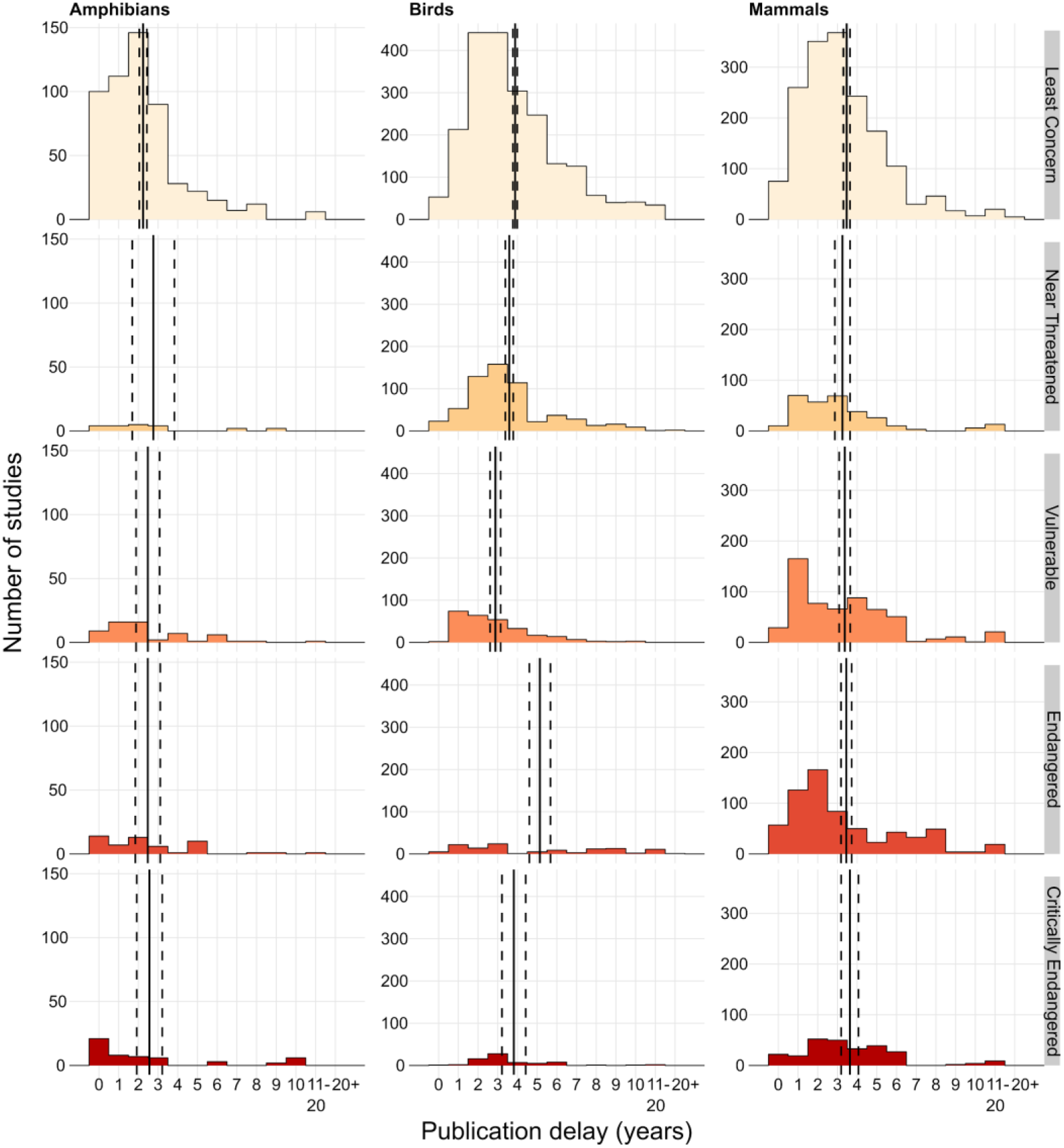
Publication delay of studies of conservation interventions (in years) grouped by the IUCN Red List Category of the species that were studied for Amphibians (Amphibia from the Amphibian Conservation synopsis), Birds (Aves from the Bird Conservation synopsis), or Mammals (Mammalia from the Bat Conservation, Primate Conservation, and Terrestrial Mammal Conservation synopses). IUCN threatened categories include Vulnerable, Endangered, and Critically Endangered, whilst non-threatened categories include Least Concern and Near Threatened (following IUCN Red List; 2020). We did not include studies on Data Deficient and Extinct in the Wild species as there were too few studies (see Methods). Vertical solid lines show mean publication delay and dashed lines show 95% Confidence Intervals. For each taxonomic group (Amphibians, Birds, and Mammals), summary estimates were obtained from a quasi-Poisson Generalised Linear Model (GLM) with only IUCN Red List status as a fixed effect (using only data on studies from the appropriate synopses).

Overall, when pooling studies testing interventions on amphibians, birds, or mammals (Fig.S2), publication delay, there were inconsistent, but significant differences between IUCN Red List status categories: studies on Least Concern species had a significantly smaller delay than Critically Endangered species (t=3.7; p=0.002; Table S9); studies on Near Threatened species had a significantly smaller delay than Endangered and Critically Endangered species (p<0.01; Table S9); and studies on Vulnerable species had a significantly smaller delay than Least Concern, Endangered, and Critically Endangered species (p<0.01; Table S9).

When only considering Amphibians or Mammals, there were no significant differences in the mean delay of studies for different IUCN Red List categories (p>0.05; Table S9). For birds, studies on Least Concern species had a significantly smaller delay than Endangered species (t= 5.2; p<0.001; Table S9 — although not compared to other categories; p>0.05; Table S9), whilst studies on Vulnerable species had a significantly smaller delay compared to all other categories (p<0.01; Table S9), and studies on Critically Endangered species had a significantly shorter delay than for Endangered species (t=2.7; p=0.043; Table S9). Studies on Near Threatened species also had a significantly smaller delay than for Endangered species (t=6.1; p<0.001; Table S9).

## Discussion

Our results suggest that conservation decision-makers must typically wait, on average, 3.5 years for the latest evidence testing the effectiveness of conservation interventions to be published. There were significant differences in publication delay between conservation subjects (synopses) and between publication sources, where studies testing interventions on captive animals and studies from non-journal literature had a smaller delay. Although we found publication delay has marginally increased over time (1912-2020), sensitivity analyses suggested this change was weak post-1980s and there is little evidence for any substantial changes over time. Publication delay also varied inconsistently between studies on species with different IUCN Red List statuses and there was little evidence that studies on more threatened species were subject to a smaller delay.

Our results concur with previous analyses of publication delay in the wider conservation literature (ca. three years; O’Donnell et al. 2010) and similar trends found in other mission-driven disciplines. For example, studies have shown a destination journal delay of ca. 9.5 months in biomedicine (Björk & Solomon 2013), ca. ten months between the release of a press statement of trial results and publication in oncology (Qunaj et al. 2018), and that only 53% of vaccine trials were published within three years after trial completion (Manzoli et al. 2014).

In conservation, a great deal can happen in 3.5 years – a species’ population may drastically decline, new threats may emerge – and conservationists may have to take rapid action to avert biodiversity and habitat loss. Without faster access to evidence on effectiveness, there is a risk that practitioners pursue ineffective practices and mis-allocate conservation resources at a time when we cannot afford to do so. Our findings are particularly concerning given that we used a conservative approach to estimate publication delay by coarsely quantifying publication delay using differences between years (which rounds down publication delay to zero for any studies completed and published within a calendar year).

We did identify, however, that studies on captive animal interventions had a significantly smaller mean delay (ca. 2 years) compared to other conservation subjects (synopses). Possible explanations for this could be that captive animals are more easily controlled, with studies often targeting smaller numbers of individuals, under experimental conditions that are easier to plan, conduct, and write-up. It may also be that authors target a relatively narrow pool of specialist journals for publication. This could mean that rejection and resubmission to another journal, for reasons other than the quality of science, is less common. It is also possible that specific journals focused on captive animals have faster publication times. Ultimately, it is likely that a combination of reduced time to submission and faster journal publication processes have led to this smaller delay.

To better understand and minimise publication delay in scientific journals, it is useful to tease apart the potential sources of delay, namely: (1) ‘write-up delay’ (the time taken from finishing data collection to submitting a study to the first journal); (2) ‘resubmission delay’ (the time taken from submitting to the first journal to submitting to the destination journal); and (3) ‘destination journal delay’ (time taken from submitting to the destination journal to the publication of the study; see Fig.6).

**Figure 6.**
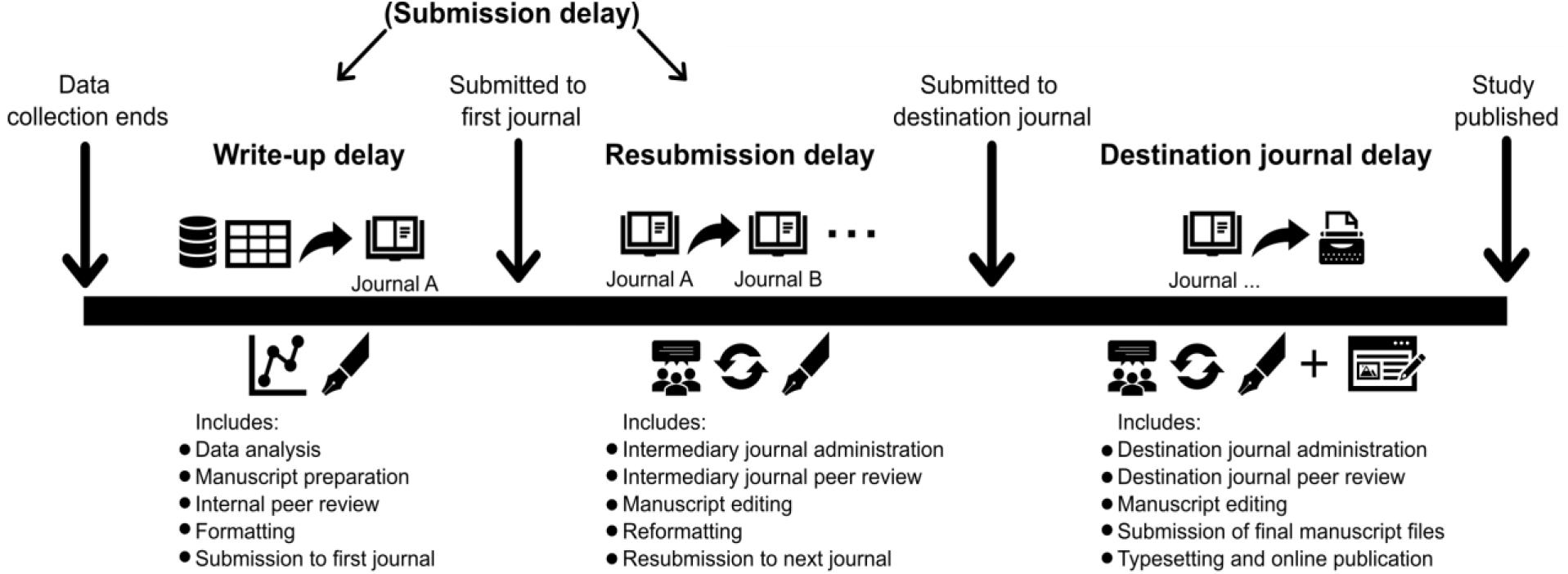
Typical publication timeline defining publication delay for studies submitted to journals and categorising different types of delay. Journals A and B are considered intermediary journals prior to being submitted to the destination journal where the study will be accepted and published. Write-up delay and resubmission delay are often combined and known collectively as ‘submission delay’ in studies investigating publication delay. Studies published in the non-journal literature would mainly suffer from write-up delay, as such studies are typically not peer-reviewed and instead progress straight to publication (sometimes after some administrative and editorial processes).

Delays in the publishing system (inc. destination journal and resubmission delay) have received much attention, with calls to speed up the review and publishing of papers in conservation (e.g. Meffe 2001; Whitten et al. 2001; Kareiva et al. 2002). Many journals have now worked to reduce processing times, and increase the efficiency of peer-review, by reducing unnecessary requirements for authors and making final manuscripts available early online (e.g., ‘early view’ prior to being published in an issue). As the publication years used in our study typically describe when studies were published in issue, this may mean we overestimated publication delay for a small number of studies. However, it has been found in other disciplines (urology and nephrology) that early view articles in 2014 were, on average, published only 95 days earlier than the final published article (Echeverría et al. 2017) – a similar delay of less than 3 months is unlikely to have substantially altered our main findings given we used differences between years as our metric of delay.

When comparing results from studies in 2002 and 2007, O’Donnell et al (2010) showed destination journal delay had reduced significantly in conservation from 572 days to 402 days, a faster decrease than in other fields they studied — although this delay is still substantial, and will hamper effective and timely conservation action. Whilst we did not find any significant decreases in publication delay over the time periods we analysed (including sensitivity analyses on more recent decades), the greater likelihood for more recent studies to have a longer delay could have masked progress in reducing publication delay to some extent — although we would argue it is unlikely that publication delay has decreased substantially, if at all. Furthermore, we found a significantly longer overall publication delay for studies published in journals (mean of ~2.7 years for recognised journals and ~2.7 for unrecognised journals) than in the non-journal literature (mean of ~2.2 years), suggesting that features typically associated with journal publication, such as peer review and editorial processes, are still contributing substantially to publication delay. Therefore, further improvements to the peer-review and publication process could reduce some of the delay we have observed.

A more systemic problem, however, is likely to be the combination of write-up delay and resubmission delay, which are collectively known as ‘submission delay’ (Fig. 1). O’Donnell et al. (2010) found a median submission delay of 696 days (1.91 years), higher than the 402 days of destination journal delay observed. Suggested reasons for submission delay can be split into: i) a lack of time, resources, or incentives to write up manuscripts in the conservation community; and ii) the time-consuming nature of the preparation, formatting, referencing, peer review, and resubmission of manuscripts. Anecdotally and from our own experiences, we also suggest that this serious problem in conservation extends to the loss of many potential papers that never complete the stage of write up, let alone submission or acceptance (i.e., ‘infinite publication delay’). An increasing length of submission delay often increases the effort and time investment needed to bring the text up-to-date and thus makes work more likely to remain unpublished.

Authors publishing studies of conservation interventions tend to be either conservation scientists in academia and conservation organisations or conservation practitioners who have tested interventions as part of their projects. When discussing the need for timely scientific contributions, Meffe (2001) suggested that “those with talents in and value to this field are seriously overcommitted”. Academics have to split their time between teaching, grant-writing, research projects, tutoring etc. (Meffe 2001). Practitioners are often juggling multiple conservation projects with limited funding, little or no time allocated to writing-up and publishing results, and limited incentives as other conservation priorities sit higher up on their agenda (O’Donnell et al. 2010). In both academia and conservation practice, short-term contracts and job insecurity can exacerbate the above and lead to rapidly changing focuses and priorities – meaning that publishing results often falls to the bottom of the pile.

At the same time, writing-up and publishing studies of interventions is not easy. Even after write-up, a manuscript may be rejected from several journals, including for subjective reasons of the perceived level of interest from readers rather than the strength of results or their importance for conservation. Substantial edits are then required to suit different journals’ formats, and reviews may suggest major changes which take time and resources to implement. It is common for published manuscripts to have gone through an iterative process of rejection and resubmission to different journals, each of which may review submissions for long time periods, leading to long resubmission delay (Vosshall 2012; Powell 2016). Indeed, a survey of 60 ecological journals showed journal rejection rates varied between 20-80%, and increased with impact factor indicating that many studies will go through multiple submission processes (Aarssen et al. 2008).

Since our quantification of publication delay for studies in the non-journal literature effectively represents write-up delay (at least to a great extent, as these studies are not typically peer-reviewed; see Fig.1), we can argue that resubmission delay generally makes up a substantial part of overall publication delay and may typically be greater than write-up delay (given that non-journal studies had a significantly smaller mean delay of ~2.2 years compared to ~3.7 years for recognised journals). Previous studies have included this resubmission delay within submission delay (see Fig. 6), but our findings tentatively suggest that write-up delay is generally smaller than the combination of resubmission delay and destination journal delay. We suggest future work could build on our results by directly quantifying and disentangling the components of publication delay observed, to help target action in areas that require more focus.

In Table 1, we present a set of possible solutions that could help to reduce write-up, resubmission, and destination journal delay. Whilst the solutions outlined in Table 1 are focussed specifically on conservation science, we believe they are relevant to many different disciplines tackling publication delay. The COVID-19 pandemic has seen a far-reaching response from the scientific community to boost the rate at which scientific research is being conducted and published (including studies of healthcare interventions) through clear incentives to publish, rapid peer-review, and streamlined editorial processes (Horbach 2020).

We believe that the conservation community could learn from this effort to build a strong evidence base that is rapidly updated with the latest studies of conservation interventions to help address the biodiversity crisis. Nevertheless, there is concern over the unavoidable trade-off between speed and quality in the dissemination of scientific evidence; for example, pre-print articles may make studies instantly accessible to decision-makers, but without rigorous peer-review, a cornerstone of the scientific publication process, such articles may contain poor quality data and analyses, and make unsubstantiated claims that are not supported by data. Accelerated publication of studies related to COVID-19 has been associated with a decline in methodological quality (Jung et al. 2021) and many retracted, disputed or heavily criticised papers (https://retractionwatch.com/retracted-coronavirus-covid-19-papers/). It is therefore crucial to minimise publication delay at each stage of the process, but not at the cost of reduced scientific rigour which may lead to poor quality evidence-based advice and ultimately ineffective, inefficient, or even harmful action.

Comprehensive and timely access to scientific evidence is vital for effective evidence-based decision making in any mission-driven discipline, but particularly in biodiversity conservation given the need to reverse the dramatic loss of biodiversity. Concerted action is required to streamline the rigorous testing and reporting of conservation interventions’ effectiveness to cover known gaps and biases in the evidence base (Christie et al. 2020, 2021). We believe our study clearly demonstrates the need for academics, practitioners, journals, organisations, and funders to work together as a scientific community to reduce publication delay as much as possible.

**Box 1.** Possible solutions to reduce publication delay for studies of conservation actions. These are only possible solutions and should not be taken as recommendations — for example, there are concerns over the dissemination of non-peer-reviewed scientific results from preprint servers and we do not advocate circumventing the peer-review process to reduce publication delay because of the risk of misinforming decision-makers.

| Component of Delay | Conservation scientists and practitioners | Journals and publishers | Funders and organisations |
| --- | --- | --- | --- |
| Write-up delay | • **Build publication into project plans.**<br>• **Collaboration between scientists and practitioners** to design experiments and publish results. Research organisations sometimes have time and money to write up and publish results.<br>• **Buddy schemes** match up individuals with others with time and knowledge suitable for analysing and writing up results.<br>• **Authors pre-register study designs and/or analyses before undertaking data collection where possible (or submit a registered report).** The extra planning involved (prior to data collection) could help reduce the need to correct errors later on or undertake more time-consuming analyses (Parker et al. 2019). | • **Less strict formatting** and structure requirements for initial submission.<br>• **Journals agree a set of styles** to reduce the cost of resubmission.<br>• **Journals produce article templates** (e.g., as for Conservation Evidence Journal and Oryx).<br>• **Journals produce article types better suited for the rapid publication of tests of interventions** (e.g., Research Notes, ‘Evidence’ articles in Conservation Science and Practice, Conservation Evidence Journal).<br>• **Offer pre-registration or publication of registered reports** to help speed up analyses and write-up when authors finish agreed methods of data collection (Parker et al. 2019). | • **Include funding, and time for writing up, in budgets.**<br>• **Include and accept published papers as project outcomes instead of reports.**<br>• **Build a culture that values the creation of evidence-base and timely dissemination of results** through training in evidence-based methods, and scientific write up.<br>• **Develop and/or use clear, standardised guidelines for writing up scientific articles.** |
| Resubmission delay | • **Authors “calibrate” submissions** to journals best suited to their work (Vosshall 2012) to avoid lengthy rejections and resubmissions.<br>• **Authors publish pre-prints online (e.g., BioRxiv, EcoEvoRxiv, SocRxiv) when the work has been submitted** to a journal. However, caution should be taken if disseminating results due to the lack of peer-review.<br>• **Authors pre-register study designs and/or analyses before undertaking data collection where possible (or submit a registered report)**. This could help reduce the likelihood of rejection and the need for lengthy resubmissions and revisions due to poor quality study design or analyses (Parker et al. 2019).<br>• **Adoption of one submission models** (e.g., ‘Peerage of Science’ and ‘Peer Community In’) that provide peer-review that multiple publishers can access, and link papers with interested journals who | • **Reduce unnecessary effort required for initial submissions** — e.g., universal formatting, word counts, flexibility in section layouts, pre-submission enquiries etc.<br>• **Reduce time spent in unnecessary rounds of review** through quick rejections, and decisive editorial decisions.<br>• **Incentivise peer review** e.g., payments or free subscription, awards for fast, high-quality reviews (Nguyen et al. 2015), or giving reviewer’s their own DOI (if reviews are transparent; Stern & O’Shea 2019).<br>• **Consider how deadlines given to peer reviewers and editors affect publication delay** and adjust these deadlines if necessary.<br>• **Consider consulting a wider pool of reviewers**, and training graduate students in peer review (Nguyen et al. 2015).<br>• **Offer pre-registration or publication of registered reports** to help the likelihood of rejection and the need for lengthy resubmissions and revisions due to poor quality study design or analyses (Parker et al. 2019).<br>• **In time-critical cases, use preliminary peer-review before submission** where journals pre-identify referees in advance (e.g., fast-tracked |  |
|  | use other's reviews to guide their decisions. | papers in Biological Conservation; Biological Conservation 2021) and/or send drafts to reviewers pre-review to allow reviewers to prepare comments (Sutherland & Lythgoe 2020).<br>• **Once accepted, publish quickly** (e.g., early view, online publishing) to reduce the time spent in publication limbo.<br>• **Embrace new initiatives of transparent peer-review** to: share reviews between potential publishers, identify papers of interest and quickly publish the already reviewed articles.<br>• **Move to a peer-review system that is "publish first, curate second"** through strengthening and increasing the use of preprint servers, allowing open, transparent peer-review, and the development of curation journals to select those articles of interest for specific audiences (Stern & O'Shea 2019). This has been realised during the COVID-19 pandemic with the creation of RR:C19 a journal that rapidly and transparently reviews and curates pre-prints (Dhar & Brand 2020). |  |
| Destination journal delay | • **Select platforms and journals that have taken steps to reduce publication delay** in the publication and peer-review process.<br>• **Consider the use of submission models (such as ‘Peer Community In’ or ‘Octopus’)** that provide **transparent peer-review and recommendation of pre-prints** or initial submissions, but without the requirement for, although compatible with, journal publication. |  | • **Promote the use of platforms and journals that have taken steps to reduce publication delay** in the publication and peer-review process. |

## Acknowledgements

We thank all those involved in collating the Conservation Evidence database for the extensive work synthesizing the evidence base used to inform this work. We thank David Williams for helpful feedback on an original version of the manuscript, and Ashley Simpkins for helpful advice. We thank Arcadia, The David and Claudia Harding Foundation and MAVA for funding.

## Footnotes

1 https://library.wwindea.org/global-statistics/

## Notes

### Competing Interest Statement

The authors have declared no competing interest.

### Summary of Updates

Corrected author's ORCID.

