## Supporting Information for "Reducing publication delay to improve the efficiency and impact of conservation science"

**DOI:**

*XXX*

### Supporting Information

Table S1 — Recognised journals (according to SCImago (2020)) for studies in the Conservation Evidence database, analysed in the current paper.

| **Recognised journals** |
|---|
| Accident Analysis And Prevention |
| Acta Agriculturae Scandinavica — Section B Soil And Plant Science |
| Acta Chiropterologica |
| Acta Herpetologica |
| Acta Horticulturae |
| Acta Oecologica |
| Acta Phytopathologica Et Entomologica Hungarica |
| Acta Theriologica Sinica |
| African Journal Of Ecology |
| African Journal Of Herpetology |
| African Journal Of Marine Science |
| Agrarforschung Schweiz |
| Agricultural And Forest Entomology |
| Agricultural Water Management |
| Agriculture, Ecosystems And Environment |
| Agroecology And Sustainable Food Systems |
| Agroforestry Systems |

| |
|---|
| Agronomy For Sustainable Development |
| Agronomy Journal |
| Agronomy Research |
| Ambio |
| American Bee Journal |
| American Journal Of Botany |
| American Journal Of Enology And Viticulture |
| American Journal Of Potato Research |
| American Journal Of Primatology |
| American Midland Naturalist |
| Amphibia — Reptilia |
| Amphibian & Reptile Conservation |
| Animal |
| Animal Behaviour |
| Animal Biology |
| Animal Conservation |
| Animal Welfare |
| Annales Botanici Fennici |
| Annales De Limnologie |
| Annales Zoologici Fennici |

| |
|---|
| Annals Of Applied Biology |
| Annals Of Botany |
| Annals Of Forest Science |
| Anthrozoos |
| Apidologie |
| Applied Animal Behaviour Science |
| Applied Entomology And Zoology |
| Applied Soil Ecology |
| Applied Vegetation Science |
| Aquacultural Engineering |
| Aquaculture |
| Aquaculture International |
| Aquaculture Nutrition |
| Aquaculture Research |
| Aquatic Conservation: Marine And Freshwater Ecosystems |
| Aquatic Ecology |
| Aquatic Ecosystem Health & Management |
| Aquatic Invasions |
| Aquatic Living Resources |
| Archives Of Environmental Contamination And Toxicology |

| |
|---|
| Arctic, Antarctic, And Alpine Research |
| Ardea |
| Auk |
| Austral Ecology |
| Australian Forestry |
| Australian Journal Of Zoology |
| Australian Mammalogy |
| Australian Zoologist |
| Basic And Applied Ecology |
| Bioagro |
| Biocontrol Science And Technology |
| Biodiversity And Conservation |
| Biofouling |
| Biological Agriculture And Horticulture |
| Biological Bulletin |
| Biological Conservation |
| Biological Control |
| Biological Invasions |
| Biological Journal Of The Linnean Society |
| Biology And Environment |

| |
|---|
| Biology And Fertility Of Soils |
| Biology Letters |
| Biomass And Bioenergy |
| Bioresource Technology |
| Bioscience |
| Biota Neotropica |
| Biotropica |
| Bird Conservation International |
| Bird Study |
| BMC Evolutionary Biology |
| Botany |
| Brazilian Journal Of Biology |
| British Birds |
| British Wildlife |
| Bryologist |
| Bulletin Of Entomological Research |
| California Agriculture |
| California Fish And Game |
| Canadian Entomologist |
| Canadian Field-Naturalist |

| |
|---|
| Canadian Journal Of Fisheries And Aquatic Sciences |
| Canadian Journal Of Forest Research |
| Canadian Journal Of Plant Pathology |
| Canadian Journal Of Soil Science |
| Canadian Journal Of Zoology |
| Catena |
| Community Ecology |
| Comparative Biochemistry And Physiology. Part A, Molecular & Integrative Physiology |
| Comparative Medicine |
| Condor |
| Conservation Biology |
| Conservation Evidence |
| Conservation Genetics |
| Conservation Letters |
| Conservation Science Western Australia |
| Copeia |
| Coral Reefs |
| Corella |
| Crop Protection |
| Crustaceana |

| |
|---|
| Cryobiology |
| Cryo-Letters |
| Diseases Of Aquatic Organisms |
| Diversity |
| Diversity And Distributions |
| Ecography |
| Ecohealth |
| Ecological Applications |
| Ecological Economics |
| Ecological Engineering |
| Ecological Entomology |
| Ecological Indicators |
| Ecological Management & Restoration |
| Ecological Research |
| Ecological Restoration |
| Ecology |
| Ecology And Evolution |
| Ecology And Society |
| Ecology Letters |
| Ecology, Environment And Conservation |

| |
|---|
| Ecosphere |
| Ecosystems |
| Emu |
| Endangered Species Research |
| Entomologia Experimentalis Et Applicata |
| Entomologia Generalis |
| Entomologica Fennica |
| Entomologist's Gazette |
| Environmental Conservation |
| Environmental Entomology |
| Environmental Evidence |
| Environmental Management |
| Environmental Pollution |
| Environmental Science And Pollution Research |
| Environmental Toxicology And Chemistry |
| Environmental Toxicology And Pharmacology |
| Estuarine, Coastal And Shelf Science |
| Ethology Ecology And Evolution |
| European Journal Of Agronomy |
| European Journal Of Forest Research |

| |
|---|
| European Journal Of Soil Biology |
| European Journal Of Soil Science |
| European Journal Of Wildlife Research |
| Field Crops Research |
| Fire Ecology |
| Fisheries Management And Ecology |
| Fisheries Research |
| Folia Geobotanica |
| Folia Horticulturae |
| Folia Primatologica |
| Folia Zoologica |
| Forest Ecology And Management |
| Forest Science |
| Forest Systems |
| Forestry |
| Forestry Chronicle |
| Freshwater Biology |
| Frontiers In Earth Science |
| Frontiers In Ecology And Evolution |
| Frontiers In Ecology And The Environment |

| |
|---|
| Functional Ecology |
| Genesis |
| Genetics And Molecular Research |
| Geoderma |
| Global Change Biology |
| Global Ecology And Conservation |
| Grass And Forage Science |
| Grassland Science |
| Great Lakes Entomologist |
| HAYATI Journal Of Biosciences |
| Herpetologica |
| Herpetological Bulletin |
| Herpetological Conservation And Biology |
| Herpetological Journal |
| Herpetological Review |
| Hormones And Behavior |
| Hortscience |
| Horttechnology |
| Human-Wildlife Interactions |
| Hydrobiologia |

| |
|---|
| Hystrix |
| Ibis |
| ICES Journal Of Marine Science |
| Insect Conservation And Diversity |
| Insect Science |
| Insectes Sociaux |
| International Agrophysics |
| International Biodeterioration And Biodegradation |
| International Journal Of Ecology |
| International Journal Of Pest Management |
| International Journal Of Primatology |
| International Journal Of Tropical Insect Science |
| International Journal Of Wildland Fire |
| International Review Of Hydrobiology |
| International Zoo Yearbook |
| Invasive Plant Science And Management |
| ISME Journal |
| Japanese Journal Of Applied Entomology And Zoology |
| Journal For Nature Conservation |
| Journal Of Agricultural And Food Chemistry |

| |
|---|
| Journal Of Agricultural Science |
| Journal Of Agronomy And Crop Science |
| Journal Of Animal And Plant Sciences |
| Journal Of Animal Ecology |
| Journal Of Animal Science |
| Journal Of Apicultural Research |
| Journal Of Applied Animal Welfare Science |
| Journal Of Applied Ecology |
| Journal Of Applied Entomology |
| Journal Of Aquatic Plant Management |
| Journal Of Arachnology |
| Journal Of Arid Environments |
| Journal Of Avian Biology |
| Journal Of Chemical Ecology |
| Journal Of Coastal Conservation |
| Journal Of Coastal Research |
| Journal Of Ecology |
| Journal Of Economic Entomology |
| Journal Of Entomological Science |
| Journal Of Environmental Education |

| |
|---|
| Journal Of Environmental Management |
| Journal Of Environmental Planning And Management |
| Journal Of Environmental Quality |
| Journal Of Environmental Studies |
| Journal Of Ethology |
| Journal Of Experimental Botany |
| Journal Of Experimental Zoology Part A: Ecological And Integrative Physiology |
| Journal Of Field Ornithology |
| Journal Of Fish And Wildlife Management |
| Journal Of Fish Biology |
| Journal Of Fish Diseases |
| Journal Of Food, Agriculture And Environment |
| Journal Of Forest Research |
| Journal Of Forest Science |
| Journal Of Forestry Research |
| Journal Of Herpetology |
| Journal Of Hymenoptera Research |
| Journal Of Insect Conservation |
| Journal Of Insect Science |
| Journal Of Invertebrate Pathology |

| |
|---|
| Journal Of Mammalogy |
| Journal Of Molluscan Studies |
| Journal Of Natural History |
| Journal Of Nematology |
| Journal Of Ornithology |
| Journal Of Pest Science |
| Journal Of Pharmacological And Toxicological Methods |
| Journal Of Plant Diseases And Protection |
| Journal Of Raptor Research |
| Journal Of Sea Research |
| Journal Of Soils And Sediments |
| Journal Of Soils And Water Conservation |
| Journal Of Sustainable Forestry |
| Journal Of The Acoustical Society Of America |
| Journal Of The American Society For Horticultural Science |
| Journal Of The Kansas Entomological Society |
| Journal Of The Marine Biological Association Of The United Kingdom |
| Journal Of The Torrey Botanical Society |
| Journal Of Threatened Taxa |
| Journal Of Tropical Ecology |

| |
|---|
| Journal Of Vegetation Science |
| Journal Of Wildlife Diseases |
| Journal Of Wildlife Management |
| Journal Of Wildlife Rehabilitation |
| Journal Of Zoo And Wildlife Medicine |
| Journal Of Zoology |
| Kasetsart Journal — Natural Science |
| Koedoe |
| Laboratory Animals |
| Lake And Reservoir Management |
| Land Degradation & Development |
| Landscape And Urban Planning |
| Landscape Ecology |
| Lichenologist |
| Limnology |
| Limosa |
| Malayan Nature Journal |
| Mammal Research |
| Mammal Review |
| Mammal Study |

| |
|---|
| Mammalia |
| Mammalian Biology |
| Management Of Biological Invasions |
| Marine And Freshwater Research |
| Marine Ecology |
| Marine Ecology — Progress Series |
| Marine Environmental Research |
| Marine Mammal Science |
| Marine Ornithology |
| Marine Policy |
| Marine Pollution Bulletin |
| Mastozoologia Neotropical |
| Medical And Veterinary Entomology |
| Medical Mycology |
| Memoirs Of The Queensland Museum |
| Memoranda — Societatis Pro Fauna Et Flora Fennica |
| Microbial Ecology |
| Mires And Peat |
| Molecular Ecology |
| Munibe Antropologia-Arkeologia |

| |
|---|
| Mycorrhiza |
| Natur Und Landschaft |
| Natural Areas Journal |
| Nature |
| Neurobiology Of Learning And Memory |
| New Forests |
| New Zealand Journal Of Botany |
| New Zealand Journal Of Crop And Horticultural Science |
| New Zealand Journal Of Zoology |
| New Zealand Plant Protection |
| North American Journal Of Aquaculture |
| North American Journal Of Fisheries Management |
| Northeastern Naturalist |
| Northwest Science |
| Notulae Botanicae Horti Agrobotanici Cluj-Napoca |
| Nutrient Cycling In Agroecosystems |
| Oecologia |
| Ohio Journal Of Sciences |
| Oikos |
| Oncoscience |

| |
|---|
| Organic Agriculture |
| Ornis Svecica |
| Ornithologische Beobachter |
| Ornitologia Colombiana |
| Ornitologia Neotropical |
| Oryx |
| Outlook On Agriculture |
| Pachyderm |
| Pacific Conservation Biology |
| Pacific Science |
| Paddy And Water Environment |
| Pakistan Journal Of Agricultural Sciences |
| Pakistan Journal Of Biological Sciences |
| Pakistan Journal Of Scientific And Industrial Research Series B: Biological Sciences |
| Pedobiologia |
| Perspectives In Plant Ecology, Evolution And Systematics |
| Pest Management Science |
| Physiological Entomology |
| Phytoparasitica |
| Phytoprotection |

| |
|---|
| Plant And Soil |
| Plant Breeding |
| Plant Disease |
| Plant Ecology |
| Plant Ecology And Diversity |
| Plant Ecology And Evolution |
| Plant Health Progress |
| Plant Systematics And Evolution |
| Planta Daninha |
| Plos ONE |
| Polar Biology |
| Polish Journal Of Ecology |
| Population Ecology |
| Preslia |
| Primate Conservation |
| Primates |
| Proceedings Of The Indian Academy Of Sciences |
| Proceedings Of The Latvian Academy Of Sciences, Section B: Natural, Exact, And Applied Sciences |
| Proceedings Of The National Academy Of Sciences Of The United States Of America |

| |
|---|
| Proceedings Of The Royal Society B: Biological Sciences |
| Progress In Materials Science |
| Rangeland Ecology & Management |
| Renewable Agriculture And Food Systems |
| Reproduction |
| Reproduction, Fertility And Development |
| Reproductive Biology And Endocrinology |
| Research In Veterinary Science |
| Restoration Ecology |
| Revista Brasileira De Ciencia Do Solo |
| Revista Brasileira De Entomologia |
| Revista De Biologia Tropical |
| Revue d'Ecologie (La Terre Et La Vie) |
| River Research And Applications |
| Royal Society Open Science |
| Russian Journal Of Ecology |
| Salamandra |
| Scandinavian Journal Of Forest Research |
| Science |
| Science For Conservation |

| |
|---|
| Scienceasia |
| Scientia Marina |
| Scientific Reports |
| Slovak Raptor Journal |
| Sociobiology |
| Soil And Tillage Research |
| Soil Biology And Biochemistry |
| Soil Research |
| Soil Science Society Of America Journal |
| Soil Use And Management |
| South African Journal Of Botany |
| South African Journal Of Enology And Viticulture |
| South African Journal Of Plant And Soil |
| Southeastern Naturalist |
| Southwestern Naturalist |
| Studies On Neotropical Fauna And Environment |
| Sustainability Science |
| The Scientific World Journal |
| Theriogenology |
| Therya |

| |
|---|
| Transportation Research Record |
| Tropical Conservation Science |
| Tropical Ecology |
| Tuexenia |
| Turk Tarim Ve Ormancilik Dergisi/Turkish Journal Of Agriculture And Forestry |
| Urban Ecosystems |
| Ursus |
| Vaccine |
| Victorian Naturalist |
| Water Science And Technology |
| Waterbirds |
| Weed Biology And Management |
| Weed Research |
| Weed Science |
| Weed Technology |
| Western North American Naturalist |
| Wetlands |
| Wetlands Ecology And Management |
| Wildlife Biology |
| Wildlife Research |

| |
|---|
| Wildlife Society Bulletin |
| Wilson Journal Of Ornithology |
| Zeitschrift Für Angewandte Entomologie |
| Zemdirbyste |
| Zoo Biology |
| Zoological Science |
| Zoologische Garten |
| Zoology And Ecology |

Table S2 — Unrecognised journals (according to SCImago (2020)) for studies in the Conservation Evidence database, analysed in the current paper.

| **Unrecognised journals** |
| --- |
| Állattani Közlemények |
| Acta Academiae Agriculturae Ac Technicae Olstenensis, Agricultura |
| Acta Biologica Slovenica |
| Acta Botanica Neerlandica |
| Acta Hydrobiologica Sinica |
| Acta Jutlandica |
| African Primates |
| Airo |
| Alces |
| Applied Herpetology |
| Australian Journal Of Experimental Agriculture |
| Australian Journal Of Experimental Agriculture And Animal Husbandry |
| Barbastella |
| Bat Research News |
| Bee World |
| Bellbird |
| Biota |
| Botanica Helvetica |

| |
|---|
| Botanical Journal Of Scotland |
| British Journal Of Entomology And Natural History |
| Bulletin De La Societe Herpetologique De France |
| Chiroptera Neotropical |
| Dodo |
| Ecología |
| Entomophaga |
| Field Studies |
| Florida Scientist |
| Galemys |
| Gibbon Journal |
| Gibier Faune Sauvage, Game Wildlife |
| Gorilla Journal |
| Herpetofauna |
| Integrated Pest Management Reviews |
| Journal For Nature Conservation And Applied Landscape Ecology |
| Journal Of Agriculture And Social Sciences |
| Journal Of Bat Research And Conservation |
| Journal Of Herpetological Medicine And Surgery |
| Journal Of The Elisha Mitchell Scientific Society |

| |
|---|
| Journal Of The Iowa Academy Of Science |
| Journal Of Wetlands Environmental Management |
| Journal Of Zoo And Aquarium Research |
| Journal Of Zoo And Aquatic Research |
| Lemur News |
| Lutra |
| Mitteilungen Der Deutschen Gesellschaft Fuer Allgemeine Und Angewandte Entomologie |
| Oceanographic Literature Review |
| Peanut Science |
| Pianura |
| Phyton (Horn) |
| Procedia Environmental Sciences |
| RAVON |
| Schweiz. Peckiana |
| Scientific Proceedings Of The Royal Dublin Society, Series A |
| South African Journal Of Wildlife Research |
| Strix |
| Swedish Journal Of Agricultural Research |
| Tearmann |

| |
|---|
| Transactions Of The Illinois State Academy Of Science |
| Transactions Of The Missouri Academy Of Science |
| Transactions Of The Western Section Of The Wildlife Society |
| Vagos |
| Virginia Journal Of Science |
| Zeitschrift Für Jagdwissenschaft |
| Zeledonia |

Table S3 — Publication sources for the non-journal literature that contributed studies to analyses in this paper. Names are as entered in the Conservation Evidence database.

| **Non-journal literature sources** |
|---|
| 10th Annual Conference Of The International Association For Landscape Ecology |
| 10th International Congress Of Plant Protection: Plant Protection For Human Welfare, 20-25 November, 1983 |
| 130 Proceedings-Vertebrate Pest Conference |
| 15th Meeting Of The European Grassland Federation |
| 1997 Brighton Crop Protection Conference — Weeds, Conference Proceedings |
| 19th Meeting Of The European Grassland Federation |
| 2005 Annual International Research Conference On Methyl Bromide Alternatives And Emissions Reductions, 31st October-3rd November, 2005 |
| 24th Vertebrate Pest Conference |
| 7th International Safflower Conference |
| A Report Submitted To The Bats And Wind Energy Cooperative. |
| Abstracts Of The EWRS-Symposium 2007, Hamar, Norway |
| African Bird Club Bulletin |
| African Entomology Memoir |
| Agrarokologie |
| Agroforestry Forum |
| Alternative Functions Of Grassland. Proceedings Of The 15th European Grassland Federation Symposium, 7-9 September 2009 |
| American Fisheries Society Symposium |

| |
|---|
| American Zoo And Aquarium Association Annual Conference Proceedings |
| Amphibian Ark Newsletter |
| Amphibian Declines: The Conservation Status Of United States Species |
| Amphibians And Roads: Proceedings Of The Toad Tunnel Conference |
| Amphibians And Roads: Toad Tunnel Conference |
| Análisis Espacial Y Representación Geográfica: Innovación Et Aplicación |
| Anzeiger Fur Schadlingskunde |
| Arthropod Natural Enemies In Arable Land |
| Arthropod Natural Enemies In Arable Land I — Density, Spatial Heterogeneity And Dispersal, Acta Jutlandica |
| Avian Landscape Ecology: Pure And Applied Issues In The Large-Scale Ecology Of Birds |
| Bats And Forests Symposium |
| Bats In Captivity Volume 2: Aspects Of Rehabilitation |
| Biodiversity And Insect Pests: Key Issues For Sustainable Management |
| Biodiversity News |
| Boletin De Sanidad Vegetal, Plagas |
| Bollettino Dell'istituto Di Entomologia Della Università Di Bologna |
| Braunschweiger Naturkundliche Schriften |
| British Crop Protection Conference: Pests And Diseases |
| British Crop Protection Council Monographs |

| |
|---|
| British Grassland Society Fifth Research Conference |
| British Grassland Society Occasional Symposium |
| British Herpetological Society Bulletin |
| British Sugar Beet Review |
| BTO Research Report |
| Bulletin Français De La Píache Et De La Pisciculture |
| Bulletin OEPP |
| Bulletin Of The Association Of Reptilian And Amphibian Veterinarians |
| Bulletin Of The Maryland Herpetological Society |
| Bulletin Of The Virginia Water Resources Center |
| Bulletin OILB SROP |
| California Energy Commission Report |
| Captive Management And Conservation Of Amphibians And Reptiles, Contributions To Herpetology Vol. 11, Society For The Study Of Amphibians And Reptiles |
| Carabid Beetles: Ecology And Evolution |
| CEFAS Final Contract Report C5256 |
| Centre For Ecology And Hydrology Defra Project Code PH0422 |
| Centre For Evidence-Based Conservation |
| Changes In The Fauna Of Wild Bees In Europe |
| Cities And The Environment |

| |
|---|
| Coastal Meadow Management — Best Practice Guidelines |
| Collaboration For Environmental Evidence |
| Conservation And Management Of Great Crested Newts |
| Creating New Habitats In Intensively Used Farmland |
| De Levende Natuur |
| Declines And Disappearances Of Australian Frogs |
| Defra |
| Department Of Biological Sciences |
| Department Of Ecology & Evolutionary Biology |
| Desert Bighorn Council Transactions |
| Deutsche Gesellschaft Für Herpetologie Und Terrarienkunde |
| Durrell Institute Of Conservation And Ecology |
| Ecological Society Of America Annual Meeting Abstracts |
| Ecology And Conservation Of Lowland Farmland Birds. Spring Conference Of The British Ornithologists' Union, 27-28 March 1999 |
| Ecology And Integrated Farming Systems |
| Ecosystems And Sustainable Development III, Advances In Ecological Sciences |
| Effect Of Sward Type And Management On Diversity Of Upland Birds. |
| Eighth International Herpetological Symposium |
| Enact |

| |
|---|
| Endangered Species Bulletin |
| Endangered Species Update |
| Environmental Encounters Series: Workshop On Ecological Corridors For Invertebrates: Strategies Of Dispersal And Recolonisation In Today's Agricultural And Forestry Landscapes |
| EPOPS |
| European And Mediterranean Plant Protection Organization |
| European And Mediterranean Plant Protection Organization Report Number 09-15078 Rev |
| Exmoor Mires Partnership |
| Fencing For Conservation. Restriction Of Evolutionary Potential Or A Riposte To Threatening Processes? |
| Fibl Dossier |
| Field Margins: Integrating Agriculture And Conservation |
| Forage For Bees In An Agricultural Landscape |
| Forestry Commission Report |
| Forward With Grass Into Europe: British Grassland Society Winter Meeting |
| French Peatland Coordination Centre |
| Froglife Conservation Report No.1 |
| Froglog |
| Frogs In The Community |
| Global Re-Introduction Perspectives: 2008. Re-Introduction Case-Studies From Around The Globe |

| |
|---|
| Global Re-Introduction Perspectives: 2010. Additional Case Studies From Around The Globe |
| Global Re-Introduction Perspectives: 2011. More Case Studies From Around The Globe |
| Grassland Science In Europe |
| Grazing Management: The Principles And Practice Of Grazing, For Profit And Environmental Gain, Within Temperate Grassland Systems: Proceedings Of The British Grassland Society Conference, 29 February-2 March, 2000 |
| Habitat Fragmentation & Infrastructure |
| Hedgerows Of The World: Their Ecological Functions In Different Landscapes, International Association For Landscape Ecology, 10th Annual Conference Of The International Association For Landscape Ecology |
| Herpetofauna And Roads Workshop — Is There Light At The End Of The Tunnel? |
| Herpetologica Bonnensis |
| Herpetological Society Bulletin |
| High Value Grassland, British Grassland Society Occasional Symposium No.38 |
| High Value Grassland: Providing Biodiversity, A Clean Environment And Premium Products. British Grassland Society Occasional Symposium No.38 |
| High Value Grassland: Providing Biodiversity, A Clean Environment And Premium Products. . British Grassland Society Occasional Symposium No.38 |
| High Value Grassland: Providing Biodiversity, A Clean Environment And Premium Products, British Grassland Society Occasional Symposium |
| How To Protect Or What We Know About Carabid Beetles: From Knowledge To Application, From Wijster (1969) To Tuczno (2001), 2002 Conference |
| ICROFS News |
| IFOAM 2000: The World Grows Organic |

| |
|---|
| In Practice: Bulletin Of The Chartered Institute Of Ecology And Environmental Management |
| USDA-ARS-SABCL (South American Biological Control Laboratory). Annual Report, 2011 |
| Infos-Ctifl |
| Insect Pest Management |
| International Conference On Habitat Fragmentation Due To Transportation Infrastructure |
| International Occasional Symposium Of The European Grassland Federation. |
| International Symposium And Workshop On Tropical Peatland |
| IOBC/Wprs Bulletin |
| IOBC/WPRS Bulletin |
| ITE Symposium |
| Joint Meeting Between The British Grassland Society And The British Ecological Society: Grassland Management And Nature Conservation. British Grassland Society Occasional Symposium |
| Kalimantan Forests And Climate Partnership |
| Land Retirement Demonstration Project Five Year Report |
| Landscape Management For Functional Biodiversity 2nd Working Group Meeting. 16-19 May 2006. |
| Landscape Management For Functional Biodiversity, 2nd Working Group Meeting |
| LIFE Project: Anglesey & Llyn Fens |
| Livestock Farming Systems: Integrating Animal Science Advances In The Search Of Sustainability |

| |
|---|
| London Naturalist |
| Möglichkeiten Und Grenzen Der Ökologisierung Der Landwirtschaft: Wissenschaftliche Grundlagen Und Praktische Erfahrungen; Beiträge Aus Dem Arbeitskreis |
| The humble bee: its life history and how to domesticate it. |
| Mededelingen Van De Faculteit Landbouwwetenschappen Universiteit Gent |
| Mitteilungen Der Biologischen Bundesanstalt Für Land-U. Forstwirtschaft |
| National Park Service Point Reyes National Seashore, California, USA |
| Natura Societa Italiana Di Scienze Naturale E Museo Civico Di Storia Naturale Milan |
| Natuurhistorisch Maandblad |
| Neotropical Primates |
| NERI, Technical Report |
| New Forest Plants Project, UK |
| Our Living Resources: A Report To The Nation On The Distribution, Abundance, And Health Of US Plants, Animals, And Ecosystems |
| Penny Anderson Associates Report |
| Pesticides, Cereal Farming And The Environment: The Boxworth Project |
| Phd Thesis |
| Plant Invasions: General Aspects And Special Problems |
| Plant Research International, Wageningen |
| Primate Tourism: A Tool For Conservation |
| Proceedings — Brighton Crop Protection Conference |

| |
|---|
| Proceedings 20th German Conference On Weed Biology And Weed Control |
| Proceedings Of A Symposium On Cheetahs As Game Ranch Animals |
| Proceedings Of National Avian-Wind Power Planning Meeting IV |
| Proceedings Of The 1987 International Crane Workshop International Crane Foundation, |
| Proceedings Of The 1989 American Association Of Zoological Parks And Aquariums National Conference |
| Proceedings Of The 1998 International Conference On Wildlife Ecology And Transportation |
| Proceedings Of The 1999 International Conference On Wildlife Ecology And Transportation |
| Proceedings Of The 2001 International Conference On Ecology And Transportation |
| Proceedings Of The 2003 International Conference On Ecology And Transportation |
| Proceedings Of The 2005 International Conference On Ecology And Transportation |
| Proceedings Of The 2007 International Conference On Ecology And Transportation |
| Proceedings Of The 5th Annual Reptile Symposium On Captive Propagation And Husbandry |
| Proceedings Of The Asian Wetland Symposium |
| Proceedings Of The British Grassland Society/British Ecological Society Conference |
| Proceedings Of The Conservation And Management Of Great Crested Newts |
| Proceedings Of The Eastern Wildlife Damage Control Conference |
| Proceedings Of The Eastern Wildlife Damage Management Conference |
| Proceedings Of The Eleventh Vertebrate Pest Conference |

| |
|---|
| Proceedings Of The Fourth International Workshop, 26-29 November 2001 |
| Proceedings Of The Hedgerow Management And Nature Conservation: British Ecological Society Conservation Ecology Group |
| Proceedings Of The HGCA Conference, Arable Crop Protection In The Balance: Profit And The Environment |
| Proceedings Of The International Conference On Wildlife Ecology And Transportation |
| Proceedings Of The International Symposium On Tropical Peatlands |
| Proceedings Of The International Union Of Game Biologists 17th Congress |
| Proceedings Of The Lowland Farmland Birds III: Delivering Solutions In An Uncertain World |
| Proceedings Of The Ninth Wildlife Damage Management Conference |
| Proceedings Of The Relationship Between Nature Conservation, Biodiversity And Organic Agriculture |
| Proceedings Of The Sixteenth Vertebrate Pest Conference |
| Proceedings Of The Sixth Workshop For Tropical Agricultural Entomologists, May 1998 |
| Proceedings Of The South Dakota Academy Of Science |
| Proceedings Of The Tenth Vertebrate Pest Conference |
| Proceedings Of The Thirteenth Australian Weeds Conference |
| Proceedings Of The Trends In Addressing Transportation Related Wildlife Mortality: Transportation Related Wildlife Mortality Seminar, FL-ER-58-96 |
| Proceedings Of The Trends In Addressing Wildlife Mortality: Transportation Related Wildlife Mortality Seminar, FL-ER-58-96 |
| Proceedings The Joint Meeting Between The British Grassland Society And The British Ecological Society. Grassland Management And Nature Conservation |

| |
|---|
| Proceedings, Soil And Crop Science Society Of Florida |
| Protecting Threatened Bats At Coal Mines: A Technical Interactive Forum |
| Raised Bog Management For Biological Diversity Conservation In Latvia |
| Recent Developments In Cereal Production. |
| Re-Creating Plant And Beetle Assemblages Of Species-Rich Chalk Grasslands On Ex-Arable Land |
| Regional Meetings Of The American Association Of Zoological Parks And Aquariums |
| Rehabilitation Of Oiled African Penguins: A Conservation Success Story |
| Re-Introduction News |
| RELU Policy And Practice Note, Number 37 Report |
| Rencontre Recherche Ruminants |
| Report For Confederación Hidrográfica Del Ebro |
| Report To Infra Econetwork Europe (IENE), Fifth IENE Meeting |
| Report To INRENA |
| Report To Syngenta |
| Report To The Nature Conservancy Council (GB) |
| Restoration Of Endangered Species: Conceptual Issues, Planning And Implementation |
| Restoration Of Temperate Wetlands |
| RSPB |
| School Of Graduate Studies And Research |

| |
|---|
| Scottish Natural Heritage Reports |
| Seabird Bycatch: Trends, Roadblocks And Solutions |
| Second International Symposium On Biological Control Of Arthropods, September 12-16, 2005 |
| Simposio: Estado Del Conocimiento De La Biología Reproductiva De Las Especies De Loros Amenazados De Colombia Con Énfasis En Iniciativas De Conservación |
| Simposio: Estado Del Conocimiento De La Biología Reproductiva De Las Especies De Loros Amenazados De Colombia Con Énfasis En Iniciativas De Conservación. II Congreso Colombiano De Zoología |
| Sonoma State University And Marin/Sonoma Mosquito And Vector Control District, Sonoma, California |
| Status And Conservation Of Midwestern Amphibians |
| Suffolk Natural History |
| Systematic Review No. 11. Collaboration For Environmental Evidence / Centre For Evidence-Based Conservation |
| Technical Bulletin, Agricultural Experiment Station, University Of Maine |
| Terra Australis |
| The Brighton Crop Protection Conference — Weeds |
| The Ecology And Conservation Of Lowland Farmland Birds |
| The Ecology Of Temperate Cereal Fields |
| The National Trust Conservation Newsletter |
| The Structure And Functioning Of Flower-Visiting Insect Communities On Hay Meadows |
| Timber, Tourists And Temples. Conservation And Development In The Maya Forest Of Belize, Guatemala And Mexico |

| |
|---|
| Tropical Peat Swamp Forest Silviculture In Central Kalimantan |
| TWSG News |
| Unpublished Report Commissioned By The National Trust |
| Unpublished Report To The West Sussex Heathlands Project |
| Urban Herpetology |
| Use Of The Tubingen Mix For Bee Pasture In Germany |
| Verhandlungen Der Gesellschaft Fur Okologie |
| Vingtième Conférence Du Columa Journées Internationales Sur La Lutte Contre Les Mauvaises Herbes |
| Wading Birds |
| WAIT School Of Biology Bulletin |
| Washington Sea Grant Program, University Of Washington Report |
| Western Foundation Of Vertebrate Zoology |
| Wolves: Behavior, Ecology, And Conservation |
| ZEF Bonn |

Table S4 — Mean publication delay (and associated 95% Confidence Intervals (CI) and numbers of studies) for each Conservation Evidence synopsis (a collection of studies based on the conservation subject in which interventions have been tested). Mean values and 95% CIs are presented in Figure 1 (Main Text) and were determined using a simple GLM (see Methods in Main Text) with only synopsis as a fixed effect. The number of studies for each synopsis may not match the Conservation Evidence website as the database is being dynamically updated with more studies over time. In addition, studies can be present in multiple synopses, and for the purposes of our analyses we also excluded reviews and meta-analyses, as well as studies with no end date of data collection.

| Synopsis | Mean | Upper 95% CI | Lower 95% CI | Number of studies |
|---|---|---|---|---|
| Amphibian Conservation | 2.31 | 2.52 | 2.11 | 547 |
| Bat Conservation | 2.82 | 3.17 | 2.48 | 237 |
| Bee Conservation | 3.01 | 3.42 | 2.59 | 175 |
| Bird Conservation | 3.98 | 4.14 | 3.82 | 1548 |
| Control of Freshwater Invasive Species | 2.41 | 2.86 | 1.95 | 115 |
| Farmland Conservation | 3.73 | 3.93 | 3.54 | 971 |
| Forest Conservation | 4.16 | 4.48 | 3.84 | 399 |
| Management of Captive Animals | 1.97 | 2.41 | 1.53 | 101 |
| Mediterranean Farmland | 3.70 | 3.92 | 3.48 | 737 |
| Natural Pest Control | 3.80 | 4.28 | 3.32 | 163 |
| Peatland Conservation | 3.77 | 4.14 | 3.41 | 277 |
| Primate Conservation | 2.66 | 2.94 | 2.38 | 343 |
| Shrubland and Heathland Conservation | 3.59 | 4.01 | 3.18 | 207 |
| Soil Fertility | 3.92 | 4.36 | 3.47 | 196 |
| Subtidal Benthic Invertebrate Conservation | 4.35 | 4.78 | 3.91 | 226 |
| Sustainable Aquaculture | 2.58 | 3.31 | 1.86 | 48 |
| Terrestrial Mammal Conservation | 3.72 | 3.89 | 3.54 | 1187 |

Table S5 — Results of quasi-Poisson Generalised Linear Model (GLM) (see Methods). The synopsis 'Management of Captive Animals' and publication source 'non-journal literature' were set as the intercept. p-values of 0.000 represent p<0.001.

| Parameter | Estimate | Standard Error | t-value | p-value |
|---|---|---|---|---|
| Intercept | -8.044 | 2.314 | -3.476 | 0.001 |
| Publication Delay | 0.004 | 0.001 | 3.598 | 0.000 |
| Farmland Conservation | 0.696 | 0.117 | 5.954 | 0.000 |
| Bird Conservation | 0.724 | 0.116 | 6.251 | 0.000 |
| Natural Pest Control | 0.698 | 0.131 | 5.335 | 0.000 |
| Control of Freshwater Invasive Species | 0.295 | 0.149 | 1.972 | 0.049 |
| Shrubland and Heathland Conservation | 0.580 | 0.128 | 4.525 | 0.000 |
| Terrestrial Mammal Conservation | 0.638 | 0.116 | 5.489 | 0.000 |
| Bat Conservation | 0.374 | 0.130 | 2.884 | 0.004 |
| Amphibian Conservation | 0.263 | 0.123 | 2.149 | 0.032 |
| Bee Conservation | 0.441 | 0.134 | 3.298 | 0.001 |
| Forest Conservation | 0.715 | 0.120 | 5.941 | 0.000 |
| Primate Conservation | 0.340 | 0.126 | 2.706 | 0.007 |
| Peatland Conservation | 0.680 | 0.124 | 5.473 | 0.000 |
| Mediterranean Farmland | 0.589 | 0.118 | 4.993 | 0.000 |
| Subtidal Benthic Invertebrate Conservation | 0.751 | 0.125 | 6.014 | 0.000 |
| Soil Fertility | 0.662 | 0.128 | 5.184 | 0.000 |
| Sustainable Aquaculture | 0.241 | 0.184 | 1.311 | 0.19 |
| Publication source — Recognised Journal | 0.434 | 0.044 | 9.908 | 0.000 |
| Publication source — Unrecognised Journal | 0.247 | 0.077 | 3.212 | 0.001 |

Table S6 — Results of Tukey's all-pair comparisons (using results from quasi-Poisson Generalised Linear Model — see Methods) to test for statistically significant differences between the publication delay of studies in different synopses. Significance level = 0.05. p-values of 0.000 represent p<0.001.

| Comparison | Estimate | Standard error | t-statistic | Adjusted p-value |
|---|---|---|---|---|
| Farmland Conservation — Management of Captive Animals | 0.696 | 0.117 | 5.954 | 0.000 |
| Bird Conservation — Management of Captive Animals | 0.724 | 0.116 | 6.251 | 0.000 |
| Natural Pest Control — Management of Captive Animals | 0.698 | 0.131 | 5.335 | 0.000 |
| Control of Freshwater Invasive Species — Management of Captive Animals | 0.295 | 0.149 | 1.972 | 0.830 |
| Shrubland and Heathland Conservation — Management of Captive Animals | 0.580 | 0.128 | 4.525 | 0.001 |
| Terrestrial Mammal Conservation — Management of Captive Animals | 0.638 | 0.116 | 5.489 | 0.000 |
| Bat Conservation — Management of Captive Animals | 0.374 | 0.130 | 2.884 | 0.204 |
| Amphibian Conservation — Management of Captive Animals | 0.263 | 0.123 | 2.149 | 0.716 |
| Bee Conservation — Management of Captive Animals | 0.441 | 0.134 | 3.298 | 0.066 |
| Forest Conservation — Management of Captive Animals | 0.715 | 0.120 | 5.941 | 0.000 |
| Primate Conservation — Management of Captive Animals | 0.340 | 0.126 | 2.706 | 0.304 |
| Peatland Conservation — Management of Captive Animals | 0.680 | 0.124 | 5.473 | 0.000 |

| | | | | |
|---|---|---|---|---|
| Mediterranean Farmland — Management of Captive Animals | 0.589 | 0.118 | 4.993 | 0.000 |
| Subtidal Benthic Invertebrate Conservation — Management of Captive Animals | 0.751 | 0.125 | 6.014 | 0.000 |
| Soil Fertility — Management of Captive Animals | 0.662 | 0.128 | 5.184 | 0.000 |
| Sustainable Aquaculture — Management of Captive Animals | 0.241 | 0.184 | 1.311 | 0.996 |
| Bird Conservation — Farmland Conservation | 0.028 | 0.034 | 0.818 | 1.000 |
| Natural Pest Control — Farmland Conservation | 0.002 | 0.070 | 0.029 | 1.000 |
| Control of Freshwater Invasive Species — Farmland Conservation | -0.401 | 0.100 | -4.006 | 0.006 |
| Shrubland and Heathland Conservation — Farmland Conservation | -0.117 | 0.065 | -1.797 | 0.913 |
| Terrestrial Mammal Conservation — Farmland Conservation | -0.058 | 0.036 | -1.602 | 0.967 |
| Bat Conservation — Farmland Conservation | -0.322 | 0.068 | -4.735 | 0.000 |
| Amphibian Conservation — Farmland Conservation | -0.433 | 0.053 | -8.226 | 0.000 |
| Bee Conservation — Farmland Conservation | -0.255 | 0.075 | -3.408 | 0.048 |
| Forest Conservation — Farmland Conservation | 0.019 | 0.048 | 0.401 | 1.000 |
| Primate Conservation — Farmland Conservation | -0.356 | 0.060 | -5.94 | 0.000 |
| Peatland Conservation — Farmland Conservation | -0.016 | 0.057 | -0.291 | 1.000 |
| Mediterranean Farmland — Farmland Conservation | -0.107 | 0.042 | -2.566 | 0.398 |
| Subtidal Benthic Invertebrate Conservation — Farmland Conservation | 0.055 | 0.058 | 0.934 | 1.000 |

| | | | | |
|---|---|---|---|---|
| Soil Fertility — Farmland Conservation | -0.034 | 0.064 | -0.536 | 1.000 |
| Sustainable Aquaculture — Farmland Conservation | -0.455 | 0.147 | -3.105 | 0.116 |
| Natural Pest Control — Bird Conservation | -0.026 | 0.068 | -0.382 | 1.000 |
| Control of Freshwater Invasive Species — Bird Conservation | -0.429 | 0.099 | -4.324 | 0.002 |
| Shrubland and Heathland Conservation — Bird Conservation | -0.144 | 0.063 | -2.306 | 0.597 |
| Terrestrial Mammal Conservation — Bird Conservation | -0.086 | 0.032 | -2.655 | 0.337 |
| Bat Conservation — Bird Conservation | -0.350 | 0.067 | -5.253 | 0.000 |
| Amphibian Conservation — Bird Conservation | -0.461 | 0.051 | -9.093 | 0.000 |
| Bee Conservation — Bird Conservation | -0.283 | 0.073 | -3.887 | 0.009 |
| Forest Conservation — Bird Conservation | -0.009 | 0.045 | -0.190 | 1.000 |
| Primate Conservation — Bird Conservation | -0.384 | 0.058 | -6.617 | 0.000 |
| Peatland Conservation — Bird Conservation | -0.044 | 0.055 | -0.813 | 1.000 |
| Mediterranean Farmland — Bird Conservation | -0.135 | 0.039 | -3.482 | 0.037 |
| Subtidal Benthic Invertebrate Conservation — Bird Conservation | 0.027 | 0.056 | 0.474 | 1.000 |
| Soil Fertility — Bird Conservation | -0.062 | 0.062 | -1.007 | 1.000 |
| Sustainable Aquaculture — Bird Conservation | -0.483 | 0.146 | -3.317 | 0.063 |
| Control of Freshwater Invasive Species — Natural Pest Control | -0.403 | 0.116 | -3.469 | 0.038 |

| | | | | |
|---|---|---|---|---|
| Shrubland and Heathland Conservation — Natural Pest Control | -0.119 | 0.087 | -1.356 | 0.994 |
| Terrestrial Mammal Conservation — Natural Pest Control | -0.060 | 0.069 | -0.871 | 1.000 |
| Bat Conservation — Natural Pest Control | -0.324 | 0.090 | -3.600 | 0.024 |
| Amphibian Conservation — Natural Pest Control | -0.435 | 0.079 | -5.501 | 0.000 |
| Bee Conservation — Natural Pest Control | -0.257 | 0.095 | -2.706 | 0.304 |
| Forest Conservation — Natural Pest Control | 0.017 | 0.076 | 0.228 | 1.000 |
| Primate Conservation — Natural Pest Control | -0.358 | 0.084 | -4.262 | 0.002 |
| Peatland Conservation — Natural Pest Control | -0.018 | 0.082 | -0.226 | 1.000 |
| Mediterranean Farmland — Natural Pest Control | -0.109 | 0.072 | -1.515 | 0.981 |
| Subtidal Benthic Invertebrate Conservation — Natural Pest Control | 0.053 | 0.083 | 0.635 | 1.000 |
| Soil Fertility — Natural Pest Control | -0.036 | 0.087 | -0.419 | 1.000 |
| Sustainable Aquaculture — Natural Pest Control | -0.457 | 0.158 | -2.896 | 0.198 |
| Shrubland and Heathland Conservation — Control of Freshwater Invasive Species | 0.285 | 0.113 | 2.513 | 0.438 |
| Terrestrial Mammal Conservation — Control of Freshwater Invasive Species | 0.343 | 0.100 | 3.442 | 0.042 |
| Bat Conservation — Control of Freshwater Invasive Species | 0.079 | 0.115 | 0.690 | 1.000 |
| Amphibian Conservation — Control of Freshwater Invasive Species | -0.031 | 0.106 | -0.293 | 1.000 |

| | | | | |
|---|---|---|---|---|
| Bee Conservation — Control of Freshwater Invasive Species | 0.146 | 0.120 | 1.221 | 0.998 |
| Forest Conservation — Control of Freshwater Invasive Species | 0.421 | 0.105 | 4.021 | 0.005 |
| Primate Conservation — Control of Freshwater Invasive Species | 0.045 | 0.111 | 0.408 | 1.000 |
| Peatland Conservation — Control of Freshwater Invasive Species | 0.385 | 0.109 | 3.548 | 0.030 |
| Mediterranean Farmland — Control of Freshwater Invasive Species | 0.294 | 0.102 | 2.892 | 0.201 |
| Subtidal Benthic Invertebrate Conservation — Control of Freshwater Invasive Species | 0.456 | 0.110 | 4.160 | 0.003 |
| Soil Fertility — Control of Freshwater Invasive Species | 0.367 | 0.113 | 3.251 | 0.077 |
| Sustainable Aquaculture — Control of Freshwater Invasive Species | -0.054 | 0.174 | -0.310 | 1.000 |
| Terrestrial Mammal Conservation — Shrubland and Heathland Conservation | 0.058 | 0.064 | 0.919 | 1.000 |
| Bat Conservation — Shrubland and Heathland Conservation | -0.206 | 0.086 | -2.397 | 0.528 |
| Amphibian Conservation — Shrubland and Heathland Conservation | -0.316 | 0.075 | -4.226 | 0.002 |
| Bee Conservation — Shrubland and Heathland Conservation | -0.139 | 0.091 | -1.518 | 0.980 |
| Forest Conservation — Shrubland and Heathland Conservation | 0.136 | 0.071 | 1.918 | 0.860 |
| Primate Conservation — Shrubland and Heathland Conservation | -0.240 | 0.080 | -3.002 | 0.151 |

| | | | | |
|---|---|---|---|---|
| Peatland Conservation — Shrubland and Heathland Conservation | 0.100 | 0.077 | 1.298 | 0.996 |
| Mediterranean Farmland — Shrubland and Heathland Conservation | 0.009 | 0.067 | 0.139 | 1.000 |
| Subtidal Benthic Invertebrate Conservation — Shrubland and Heathland Conservation | 0.171 | 0.078 | 2.191 | 0.686 |
| Soil Fertility — Shrubland and Heathland Conservation | 0.082 | 0.082 | 0.996 | 1.000 |
| Sustainable Aquaculture — Shrubland and Heathland Conservation | -0.339 | 0.156 | -2.178 | 0.694 |
| Bat Conservation — Terrestrial Mammal Conservation | -0.264 | 0.067 | -3.950 | 0.007 |
| Amphibian Conservation — Terrestrial Mammal Conservation | -0.375 | 0.052 | -7.230 | 0.000 |
| Bee Conservation — Terrestrial Mammal Conservation | -0.197 | 0.074 | -2.663 | 0.331 |
| Forest Conservation — Terrestrial Mammal Conservation | 0.077 | 0.046 | 1.671 | 0.952 |
| Primate Conservation — Terrestrial Mammal Conservation | -0.298 | 0.059 | -5.065 | 0.000 |
| Peatland Conservation — Terrestrial Mammal Conservation | 0.042 | 0.055 | 0.752 | 1.000 |
| Mediterranean Farmland — Terrestrial Mammal Conservation | -0.049 | 0.039 | -1.245 | 0.998 |
| Subtidal Benthic Invertebrate Conservation — Terrestrial Mammal Conservation | 0.113 | 0.057 | 1.982 | 0.825 |
| Soil Fertility — Terrestrial Mammal Conservation | 0.024 | 0.063 | 0.378 | 1.000 |
| Sustainable Aquaculture — Terrestrial Mammal Conservation | -0.397 | 0.146 | -2.720 | 0.293 |

| | | | | |
|---|---|---|---|---|
| Amphibian Conservation — Bat Conservation | -0.110 | 0.077 | -1.431 | 0.989 |
| Bee Conservation — Bat Conservation | 0.067 | 0.094 | 0.710 | 1.000 |
| Forest Conservation — Bat Conservation | 0.342 | 0.074 | 4.630 | 0.000 |
| Primate Conservation — Bat Conservation | -0.034 | 0.082 | -0.415 | 1.000 |
| Peatland Conservation — Bat Conservation | 0.306 | 0.080 | 3.844 | 0.011 |
| Mediterranean Farmland — Bat Conservation | 0.215 | 0.069 | 3.097 | 0.118 |
| Subtidal Benthic Invertebrate Conservation — Bat Conservation | 0.377 | 0.081 | 4.678 | 0.000 |
| Soil Fertility — Bat Conservation | 0.288 | 0.085 | 3.379 | 0.052 |
| Sustainable Aquaculture — Bat Conservation | -0.133 | 0.157 | -0.848 | 1.000 |
| Bee Conservation — Amphibian Conservation | 0.177 | 0.084 | 2.117 | 0.739 |
| Forest Conservation — Amphibian Conservation | 0.452 | 0.061 | 7.432 | 0.000 |
| Primate Conservation — Amphibian Conservation | 0.076 | 0.070 | 1.085 | 1.000 |
| Peatland Conservation — Amphibian Conservation | 0.416 | 0.067 | 6.183 | 0.000 |
| Mediterranean Farmland — Amphibian Conservation | 0.325 | 0.056 | 5.829 | 0.000 |
| Subtidal Benthic Invertebrate Conservation — Amphibian Conservation | 0.487 | 0.069 | 7.042 | 0.000 |
| Soil Fertility — Amphibian Conservation | 0.398 | 0.074 | 5.374 | 0.000 |
| Sustainable Aquaculture — Amphibian Conservation | -0.023 | 0.151 | -0.150 | 1.000 |
| Forest Conservation — Bee Conservation | 0.275 | 0.081 | 3.411 | 0.046 |
| Primate Conservation — Bee Conservation | -0.101 | 0.088 | -1.142 | 0.999 |

| | | | | |
|---|---|---|---|---|
| Peatland Conservation — Bee Conservation | 0.239 | 0.086 | 2.777 | 0.261 |
| Mediterranean Farmland — Bee Conservation | 0.148 | 0.077 | 1.924 | 0.856 |
| Subtidal Benthic Invertebrate Conservation — Bee Conservation | 0.310 | 0.087 | 3.556 | 0.028 |
| Soil Fertility — Bee Conservation | 0.221 | 0.091 | 2.431 | 0.502 |
| Sustainable Aquaculture — Bee Conservation | -0.200 | 0.160 | -1.248 | 0.998 |
| Primate Conservation — Forest Conservation | -0.376 | 0.067 | -5.609 | 0.000 |
| Peatland Conservation — Forest Conservation | -0.036 | 0.064 | -0.562 | 1.000 |
| Mediterranean Farmland — Forest Conservation | -0.127 | 0.050 | -2.529 | 0.426 |
| Subtidal Benthic Invertebrate Conservation — Forest Conservation | 0.035 | 0.065 | 0.547 | 1.000 |
| Soil Fertility — Forest Conservation | -0.054 | 0.070 | -0.766 | 1.000 |
| Sustainable Aquaculture — Forest Conservation | -0.475 | 0.149 | -3.179 | 0.094 |
| Peatland Conservation — Primate Conservation | 0.34 | 0.073 | 4.634 | 0.000 |
| Mediterranean Farmland — Primate Conservation | 0.249 | 0.062 | 3.987 | 0.006 |
| Subtidal Benthic Invertebrate Conservation — Primate Conservation | 0.411 | 0.075 | 5.505 | 0.000 |
| Soil Fertility — Primate Conservation | 0.322 | 0.079 | 4.063 | 0.004 |
| Sustainable Aquaculture — Primate Conservation | -0.099 | 0.154 | -0.644 | 1.000 |
| Mediterranean Farmland — Peatland Conservation | -0.091 | 0.059 | -1.549 | 0.976 |
| Subtidal Benthic Invertebrate Conservation — Peatland Conservation | 0.071 | 0.071 | 0.994 | 1.000 |

| | | | | |
|---|---|---|---|---|
| Soil Fertility — Peatland Conservation | -0.018 | 0.076 | -0.235 | 1.000 |
| Sustainable Aquaculture — Peatland Conservation | -0.439 | 0.152 | -2.880 | 0.207 |
| Subtidal Benthic Invertebrate Conservation — Mediterranean Farmland | 0.162 | 0.060 | 2.713 | 0.300 |
| Soil Fertility — Mediterranean Farmland | 0.073 | 0.066 | 1.109 | 0.999 |
| Sustainable Aquaculture — Mediterranean Farmland | -0.348 | 0.147 | -2.363 | 0.552 |
| Soil Fertility — Subtidal Benthic Invertebrate Conservation | -0.089 | 0.077 | -1.150 | 0.999 |
| Sustainable Aquaculture — Subtidal Benthic Invertebrate Conservation | -0.510 | 0.153 | -3.336 | 0.059 |
| Sustainable Aquaculture — Soil Fertility | -0.421 | 0.155 | -2.712 | 0.300 |

Table S7 — Results of Tukey's all-pair comparisons (using results from quasi-Poisson Generalised Linear Model — see Methods) to test for statistically significant differences between the publication delay of studies in different publication sources non-journal literature, recognised journals, and unrecognised journals according to SCImago (2020)). Significance level = 0.05. p-values of 0.000 represent p<0.001.

| Publication source comparison | Estimate | Standard error | t-statistic | Adjusted p-value |
|---|---|---|---|---|
| Recognised journals — Non-journal literature | 0.434 | 0.044 | 9.908 | 0.000 |
| Unrecognised journals — Non-journal literature | 0.247 | 0.077 | 3.212 | 0.003 |
| Unrecognised journals — Recognised journals | -0.187 | 0.066 | -2.840 | 0.012 |

Table S8 — Results of sensitivity analysis using quasi-Poisson Generalised Linear Models (GLM) for different time periods to assess whether the fact that more recently published studies are more likely to have a longer delay may have affected our results (see Methods). The synopsis ‘Management of Captive Animals’ and publication source ‘non-journal literature’ were set as the intercept. Values of 0.000 represent values less than 0.001.

| 1980-2020 | | | | |
|---|---|---|---|---|
| Parameter | Estimate | Standard Error | t-value | p-value |
| Intercept | -5.893 | 2.772 | -2.126 | 0.034 |
| Publication Delay | 0.003 | 0.001 | 2.224 | 0.026 |
| Farmland Conservation | 0.693 | 0.117 | 5.920 | 0.000 |
| Bird Conservation | 0.721 | 0.116 | 6.222 | 0.000 |
| Natural Pest Control | 0.693 | 0.131 | 5.283 | 0.000 |
| Control of Freshwater Invasive Species | 0.305 | 0.149 | 2.044 | 0.041 |
| Shrubland and Heathland Conservation | 0.574 | 0.128 | 4.476 | 0.000 |
| Terrestrial Mammal Conservation | 0.648 | 0.116 | 5.573 | 0.000 |
| Bat Conservation | 0.384 | 0.130 | 2.959 | 0.003 |
| Amphibian Conservation | 0.265 | 0.123 | 2.16 | 0.031 |
| Bee Conservation | 0.424 | 0.135 | 3.139 | 0.002 |

| | | | | |
|---|---|---|---|---|
| Forest Conservation | 0.719 | 0.120 | 5.967 | 0.000 |
| Primate Conservation | 0.339 | 0.126 | 2.699 | 0.007 |
| Peatland Conservation | 0.679 | 0.124 | 5.456 | 0.000 |
| Mediterranean Farmland | 0.593 | 0.118 | 5.024 | 0.000 |
| Subtidal Benthic Invertebrate Conservation | 0.755 | 0.125 | 6.043 | 0.000 |
| Soil Fertility | 0.672 | 0.128 | 5.254 | 0.000 |
| Sustainable Aquaculture | 0.241 | 0.184 | 1.314 | 0.189 |
| Publication source — Recognised Journal | 0.437 | 0.044 | 9.949 | 0.000 |
| Publication source — Unrecognised Journal | 0.245 | 0.077 | 3.178 | 0.001 |
| 1990-2020 | | | | |
| Intercept | -0.701 | 3.512 | -0.200 | 0.842 |
| Publication Delay | 0.000 | 0.002 | 0.270 | 0.787 |
| Farmland Conservation | 0.687 | 0.118 | 5.806 | 0.000 |
| Bird Conservation | 0.705 | 0.117 | 6.002 | 0.000 |

| | | | | |
|---|---|---|---|---|
| Natural Pest Control | 0.674 | 0.134 | 5.028 | 0.000 |
| Control of Freshwater Invasive Species | 0.312 | 0.151 | 2.060 | 0.039 |
| Shrubland and Heathland Conservation | 0.575 | 0.130 | 4.416 | 0.000 |
| Terrestrial Mammal Conservation | 0.659 | 0.118 | 5.592 | 0.000 |
| Bat Conservation | 0.405 | 0.131 | 3.081 | 0.002 |
| Amphibian Conservation | 0.280 | 0.124 | 2.257 | 0.024 |
| Bee Conservation | 0.421 | 0.136 | 3.092 | 0.002 |
| Forest Conservation | 0.719 | 0.122 | 5.905 | 0.000 |
| Primate Conservation | 0.422 | 0.127 | 3.326 | 0.001 |
| Peatland Conservation | 0.684 | 0.126 | 5.431 | 0.000 |
| Mediterranean Farmland | 0.600 | 0.119 | 5.026 | 0.000 |
| Subtidal Benthic Invertebrate Conservation | 0.762 | 0.126 | 6.037 | 0.000 |
| Soil Fertility | 0.681 | 0.129 | 5.272 | 0.000 |
| Sustainable Aquaculture | 0.238 | 0.185 | 1.292 | 0.197 |

| | | | | |
|---|---|---|---|---|
| Publication source — Recognised Journal | 0.462 | 0.045 | 10.184 | 0.000 |
| Publication source — Unrecognised Journal | 0.203 | 0.081 | 2.502 | 0.012 |
| 2000-2020 | | | | |
| Intercept | 6.687 | 4.910 | 1.362 | 0.173 |
| Publication Delay | -0.003 | 0.002 | -1.303 | 0.192 |
| Farmland Conservation | 0.652 | 0.127 | 5.143 | 0.000 |
| Bird Conservation | 0.690 | 0.126 | 5.480 | 0.000 |
| Natural Pest Control | 0.718 | 0.142 | 5.061 | 0.000 |
| Control of Freshwater Invasive Species | 0.167 | 0.158 | 1.055 | 0.291 |
| Shrubland and Heathland Conservation | 0.616 | 0.136 | 4.531 | 0.000 |
| Terrestrial Mammal Conservation | 0.638 | 0.125 | 5.083 | 0.000 |
| Bat Conservation | 0.400 | 0.135 | 2.961 | 0.003 |
| Amphibian Conservation | 0.326 | 0.131 | 2.486 | 0.013 |
| Bee Conservation | 0.365 | 0.143 | 2.552 | 0.011 |

| | | | | |
|---|---|---|---|---|
| Forest Conservation | 0.739 | 0.128 | 5.755 | 0.000 |
| Primate Conservation | 0.490 | 0.133 | 3.687 | 0.000 |
| Peatland Conservation | 0.701 | 0.132 | 5.313 | 0.000 |
| Mediterranean Farmland | 0.616 | 0.126 | 4.884 | 0.000 |
| Subtidal Benthic Invertebrate Conservation | 0.770 | 0.131 | 5.873 | 0.000 |
| Soil Fertility | 0.694 | 0.135 | 5.142 | 0.000 |
| Sustainable Aquaculture | 0.253 | 0.178 | 1.420 | 0.156 |
| Publication source — Recognised Journal | 0.416 | 0.045 | 9.276 | 0.000 |
| Publication source — Unrecognised Journal | 0.266 | 0.082 | 3.243 | 0.001 |

Table S9 — Results of Tukey's all-pair comparisons (using results from quasi-Poisson Generalised Linear Model — see Methods) to test for statistically significant differences between the publication delay of studies testing interventions on species with different IUCN Red List Status categories. IUCN Red List Status acronyms: LC = Least Concern, VU = Vulnerable, EN = Endangered, CR = Critically Endangered, EW = Extinct in the Wild. Significance level = 0.05. p-values of 0.000 represent p<0.001.

| IUCN Category comparison | Estimate | Standard error | t-statistic | Adjusted p-value |
|---|---|---|---|---|
| All | | | | |
| NT — LC | -0.065 | 0.032 | -2.048 | 0.229 |
| VU — LC | -0.110 | 0.034 | -3.284 | 0.008 |
| EN — LC | 0.079 | 0.034 | 2.334 | 0.125 |
| CR — LC | 0.191 | 0.051 | 3.719 | 0.002 |
| VU — NT | -0.046 | 0.042 | -1.079 | 0.806 |
| EN — NT | 0.144 | 0.043 | 3.352 | 0.007 |
| CR — NT | 0.256 | 0.058 | 4.436 | 0.000 |
| EN — VU | 0.189 | 0.043 | 4.408 | 0.000 |
| CR — VU | 0.302 | 0.058 | 5.179 | 0.000 |
| CR — EN | 0.112 | 0.055 | 2.026 | 0.239 |
| Amphibians | | | | |
| NT — LC | 0.185 | 0.199 | 0.930 | 0.876 |
| VU — LC | 0.033 | 0.131 | 0.249 | 0.999 |
| EN — LC | -0.001 | 0.137 | -0.010 | 1.000 |

| | | | | |
|---|---|---|---|---|
| CR — LC | 0.184 | 0.140 | 1.318 | 0.661 |
| VU — NT | -0.152 | 0.231 | -0.66 | 0.961 |
| EN — NT | -0.186 | 0.232 | -0.803 | 0.923 |
| CR — NT | -0.001 | 0.233 | -0.004 | 1.000 |
| EN — VU | -0.034 | 0.179 | -0.191 | 1.000 |
| CR — VU | 0.151 | 0.181 | 0.837 | 0.912 |
| CR — EN | 0.185 | 0.183 | 1.011 | 0.838 |
| Birds | | | | |
| NT — LC | -0.081 | 0.032 | -2.517 | 0.076 |
| VU — LC | -0.308 | 0.050 | -6.142 | 0.000 |
| EN — LC | 0.287 | 0.055 | 5.195 | 0.000 |
| CR — LC | 0.021 | 0.083 | 0.258 | 0.999 |
| VU — NT | -0.226 | 0.056 | -4.068 | 0.000 |
| EN — NT | 0.369 | 0.061 | 6.083 | 0.000 |
| CR — NT | 0.103 | 0.086 | 1.188 | 0.735 |
| EN — VU | 0.595 | 0.072 | 8.296 | 0.000 |
| CR — VU | 0.329 | 0.095 | 3.480 | 0.004 |
| CR — EN | -0.266 | 0.097 | -2.733 | 0.043 |
| Mammals | | | | |

| | | | | |
|---|---|---|---|---|
| NT — LC | -0.071 | 0.065 | -1.099 | 0.799 |
| VU — LC | -0.087 | 0.050 | -1.749 | 0.391 |
| EN — LC | -0.025 | 0.046 | -0.539 | 0.982 |
| CR — LC | 0.024 | 0.067 | 0.365 | 0.996 |
| VU — NT | -0.016 | 0.073 | -0.213 | 1.000 |
| EN — NT | 0.046 | 0.072 | 0.644 | 0.966 |
| CR — NT | 0.096 | 0.086 | 1.111 | 0.792 |
| EN — VU | 0.062 | 0.058 | 1.068 | 0.815 |
| CR — VU | 0.111 | 0.075 | 1.493 | 0.553 |
| CR — EN | 0.049 | 0.073 | 0.676 | 0.959 |

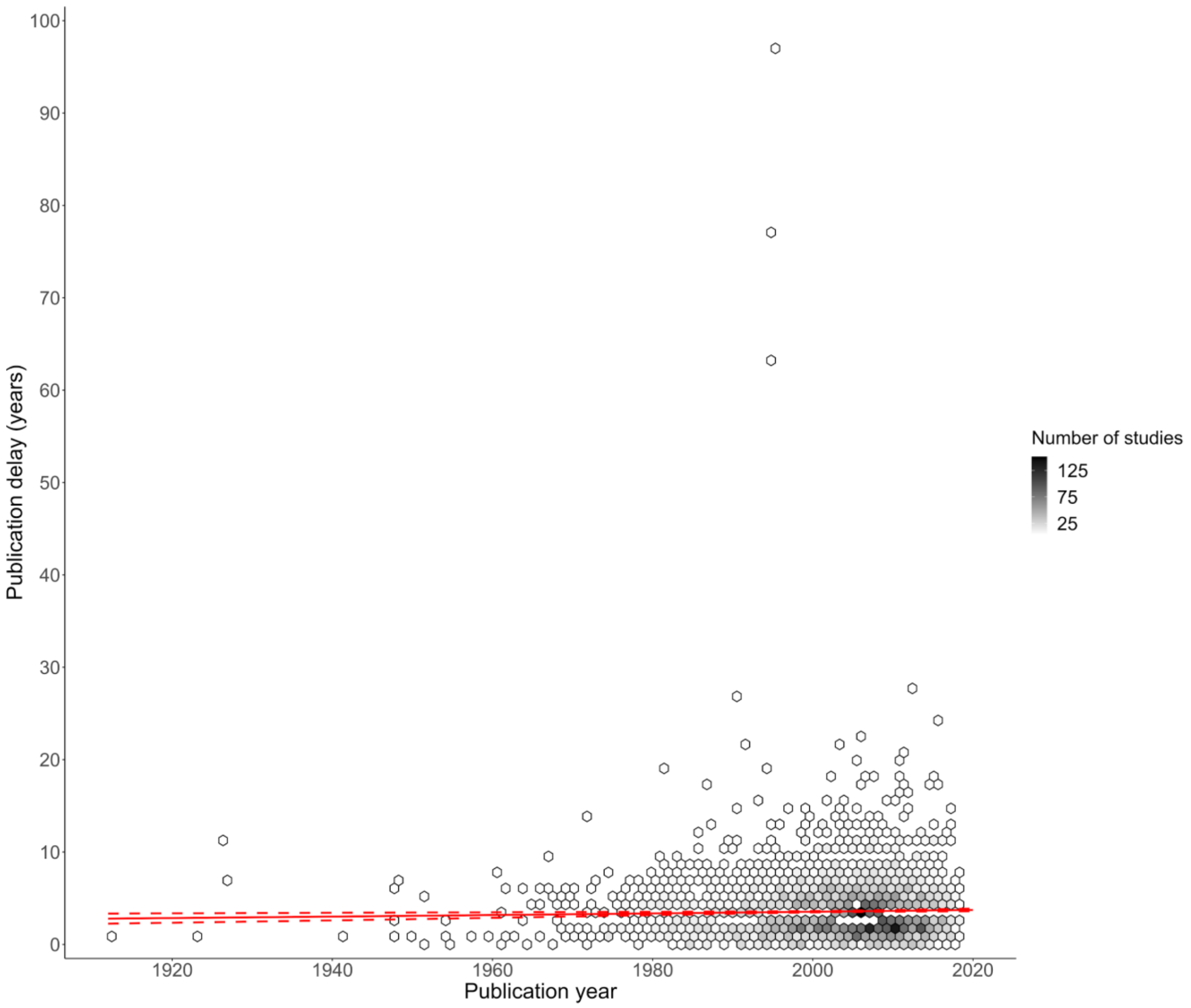


***Figure S1 – Changes in publication delay relative to the year in which studies of interventions were published. The colour of hexagons is relative to the number of data points (studies) at that position on the graph. The red solid and dotted lines represent modelled mean and 95% confidence intervals for publication delay using a quasi-Poisson Generalised Linear Model (GLM) (using only pubdate as a fixed effect — see Methods and Table S6 for full model). Data for all studies is shown here, including for a small number of studies with a publication delay greater than 20 years which were omitted from Fig.4 to improve data visualisation.***

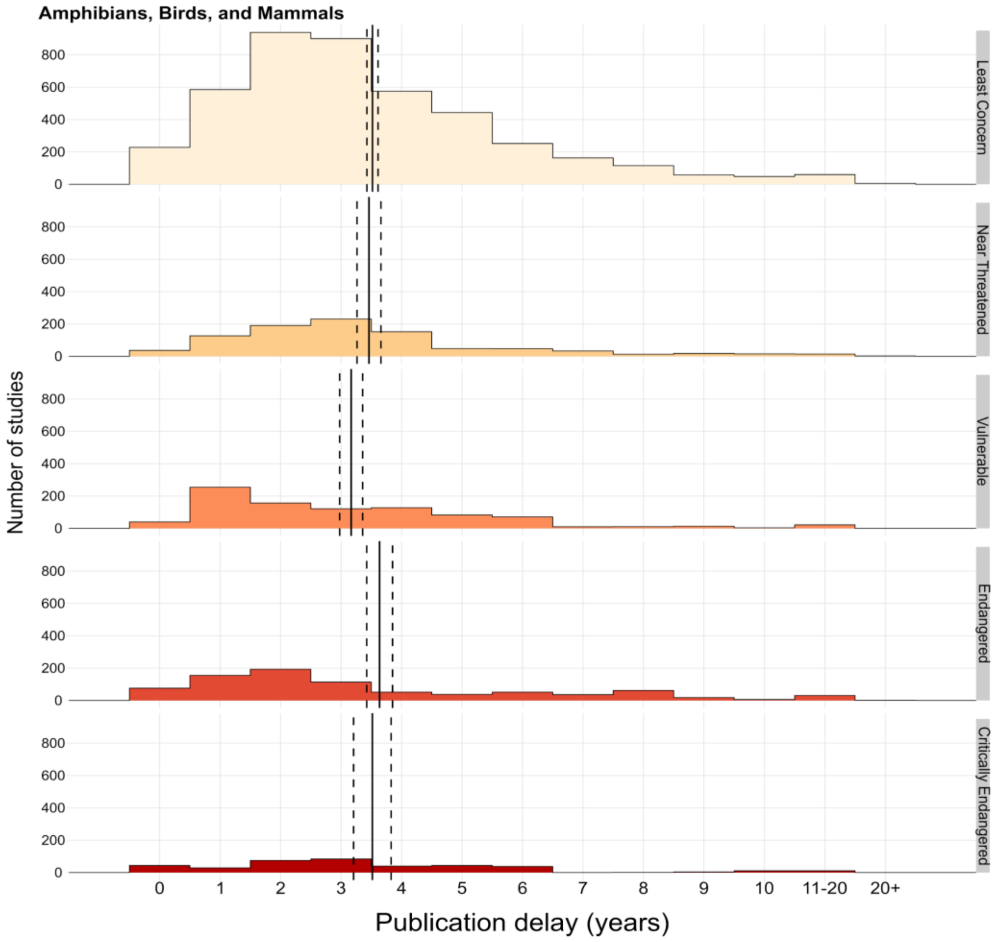


***Figure S2 – Publication delay of studies of conservation interventions (in years) grouped by the IUCN Red List Category of the species that were studied for Amphibians (Amphibia from the Amphibian Conservation synopsis), Birds (Aves from the Bird Conservation synopsis), and Mammals (Mammalia from the Bat Conservation, Primate Conservation, and Terrestrial Mammal Conservation synopses). IUCN threatened categories include Vulnerable, Endangered, and Critically Endangered, whilst non-threatened categories includes Least Concern and Near Threatened (following IUCN Red List; 2020). We did not include studies on Data Deficient and Extinct in the Wild species as there were too few studies (see Methods). Vertical solid lines show mean publication delay and dashed lines show 95% Confidence Intervals. Summary estimates were obtained from a quasi-Poisson Generalised Linear Model (GLM) with only IUCN Red List status as a fixed effect (using only data on studies from the relevant synopses above).***
